# Microtubule acetylation is required for mechanosensation in *Drosophila*

**DOI:** 10.1101/252601

**Authors:** Connie Yan, Fei Wang, Yun Peng, Claire R. Williams, Brian Jenkins, Jill Wildonger, John C. Tuthill, Yang Xiang, Stephen L. Rogers, Jay Z. Parrish

## Abstract

At the cellular level, α-tubulin acetylation alters the structure of microtubules to render them mechanically resistant to compressive forces. How this biochemical property of microtubule acetylation relates to mechanosensation remains unknown, though prior studies have shown that microtubule acetylation plays a role in touch perception. Here, we identify the major *Drosophila* α-tubulin acetylase (dTAT) and show that it plays key roles in several forms of mechanosensation while exerting little effect on other sensory modalities. dTAT is highly expressed in neurons of the larval peripheral nervous system (PNS), but is not required for normal neuronal morphogenesis. We show that mutation of the acetylase gene or the K40 acetylation site in α-tubulin impairs mechanical sensitivity in sensory neurons and behavioral responses to gentle touch, harsh touch, gravity, and sound stimulus, but not thermal stimulus. Finally, we show that dTAT is required for mechanically-induced activation of NOMPC, a microtubule-associated transient receptor potential channel, and functions to maintain integrity of the microtubule cytoskeleton in response to mechanical stimulation.

## Introduction

Mechanosensation is a signal transduction process in which mechanical forces are converted into the neuronal signals that mediate hearing, balance, proprioception, and touch. In the peripheral nervous system (PNS) this conversion is mediated by ion channels that are gated by mechanical stimuli (Coste et al., 2010; Maroto et al., 2005; O’Hagan et al., 2005; Patel et al., 1998; Walker et al., 2000). Mechanosensitive ion channels appear to have evolved multiple times and, as a result, several different channel families contribute to mechanosensation in animals, notably including transient receptor potential (TRP) channels, epithelial Na+ channel (ENaC)/degenerin (DEG) family channels, Piezos, and and transmembrane channel-like (TMC) proteins (Katta et al., 2015). Mechanical force is thought to activate the channels by inducing conformational changes, and two distinct models have been proposed to explain how force gates these channels. In the ‘force from lipids’ model the tension required for channel gating is exerted by direct interaction between the mechanoreceptor and lipids of the plasma membrane as the membrane is deformed by force (Brohawn et al., 2014; Christensen and Corey, 2007; Kung, 2005; Lewis and Grandl, 2015). In the ‘force from filaments’ model mechanical force is transduced to the channel via tethering of the channel to a non-compliant structure by a gating spring such that movement of the membrane-bound channel relative to the immobile structure induces tension within the spring leading to activation (Howard and Hudspeth, 1987; Jin et al., 2017; Liang et al., 2013). In this model, the gating spring is an elastic tether that connects the channel to extracellular structures, such as the extracellular matrix, or intracellular components, such as the cytoskeleton (Jin et al., 2017).

Much of our understanding about how tethered mechanoreceptors interact with the cytoskeleton has come from recent studies of mechanosensitive TRP channels. Notably, two TRP channel receptors, mammalian TRPV1 and *Drosophila* TRPN/NOMPC, directly interact with microtubules (Cheng et al., 2010; Prager-Khoutorsky et al., 2014). In mammalian osmosensory neurons, TRPV1 directly binds to a dense sub-cortical network of microtubules in the soma through a pair of tubulin-binding motifs in its cytoplasmic C-terminus (Prager-Khoutorsky et al., 2014). Under hypertonic conditions, cells shrink and compress their membranes against microtubules leading to an elastic compression that “pushes” the channel open. NOMPC possesses an elongated N-terminal cytoplasmic domain containing 29 tandem ankyrin repeat (AR) domains that bind to microtubules (Cheng et al., 2010; Zhang et al., 2015). Tetramers of NOMPC form a complex in which their four AR domains are organized into a quadruple bundle of helical, spring-like structures (Jin et al., 2017). The AR domains and microtubule interactions are necessary for touch-evoked responses of NOMPC, leading to the model that the AR helical elements function as the elastic tether allowing mechanical membrane deformation to gate channel opening (Howard and Bechstedt, 2004; Jin et al., 2017; Liang et al., 2013; Zhang et al., 2015). Thus, direct interaction with the microtubule cytoskeleton is necessary for TRPV1 and NOMPC to function as mechanoreceptors and hence modulating mechanical properties of microtubules could be a control point for these channels.

Post-translational modifications of microtubules regulate their function during mechanosensation in several model systems. In *C. elegans*, for example, the α-tubulin acetylase MEC-17 was identified in a screen for mutations that produced defects in mechanosensation in touch receptor neurons (Chalfie and Au, 1989). MEC-17 was the founding member of a family of α-tubulin acetylases with conserved homologues in all organisms that possess cilia (Akella et al., 2010). Depletion of zebrafish MEC-17 using morpholinos produced a variety of developmental defects in embryos including reduced startle responses to touch (Akella et al., 2010). In mice, mutants lacking the *Atat1* homologue in sensory neurons exhibited reduced mechanosensitivity and were unresponsive is assays for touch and pain (Kalebic et al., 2013a; Kim et al., 2013). These studies broadly implicate microtubules as conserved elements in mechanosensation and highlight a key regulatory role for acetylation.

Although microtubule acetylation was first discovered 30 years ago (L’Hernault and Rosenbaum, 1985), our understanding of its biological function was hindered until the recent identification of α-tubulin acetylases. Acetylation of α-tubulin occurs on lysine 40 (K40) in the lumen of the microtubule and has generally been associated with populations of long-lived microtubules. Over the last year, a series of studies have implicated α-tubulin acetylation as an adaptive biochemical mechanism that allows microtubules to resist mechanical stress. *In vitro* assays in which individual microtubules were mechanically stressed by repeated cycles of bending showed they could be damaged, resulting in decreased microtubule stiffness and localized material fatigue (Portran et al., 2017). K40 acetylation enhanced microtubule flexibility and increased their mechanical resilience by altering the structure of the microtubule lattice, allowing them to comply with deformative forces without breaking (Xu et al., 2017). In cultured cells, microtubules are compressed and broken by contractility of the actin cytoskeleton; microtubule acetylation affords them the structural resilience to resist compressive forces. If the primary role for K40 acetylation is to tune mechanical properties of microtubules, how might this biochemical activity relate to its requirement during mechanosensation?

In this study, we report the identification of *Drosophila* α-tubulin acetylase (dTAT) and our characterization of its role in mechanosensation. dTAT is broadly required for α-tubulin acetylation and is notably enriched in the PNS. We found that blocking α-tubulin acetylation broadly affected mechanosensation, but not other sensory modalities, while causing minimal effects on dendrite morphogenesis in the PNS. Using calcium imaging, we found that mutation of *dTat* or non-acetylatable alleles of *α-Tubulin84B* (*αTub84B)* attenuated gentle touch responses of NOMPC-expressing class III da (c3da) neurons, and we further found that dTAT is required for NOMPC-dependent mechanically-induced membrane depolarization. However, dTAT does not regulate gentle touch responses via effects on NOMPC-microtubule interactions or NOMPC localization. Instead, dTAT modulates mechanical stability of microtubules to control gentle touch responses and other forms of mechanosensation. First, hyperacetylation or taxol-induced microtubule stabilization sensitize larvae to gentle touch. Second, taxol treatment rescues mechanosensory behavioral defects of *dTat* mutants and non-acetylatable *αTub84B* mutants. Third, sensory neurons in *dTat* mutants accumulate microtubule breaks during development and are more sensitive to mechanically-induced microtubule breakage than wild type controls. Thus, modulation of microtubule mechanical stability appears to be a critical control point for mechanosensation.

## Results

### dTAT is the major microtubule acetylase in *Drosophila.*

Five different acetylases are capable of modifying α-tubulin in mammalian cells or *C. elegans* including GCN5, elongator protein 3 (ELP3), N-acetylase 10 (NAT10), the ARD1-NAT1 complex, and αTAT/MEC-17 (Akella et al., 2010; Conacci-Sorrell et al., 2010; Creppe et al., 2009; Ohkawa et al., 2008; Shida et al., 2010). *Drosophila* homologues for GCN5, ELP3, ARD1, and NAT1 have been studied in other contexts, but their roles in microtubule acetylation have not been characterized. We additionally identified the GNAT domain-containing lethal(1)G0020/CG1994 as the likely NAT10 homologue (53.8% identity) and CG3967 and CG17003 as potential αTAT/MEC-17 homologues (CG3967, 38.8% identity; CG17003, 34.7% identity). To test their roles in microtubule acetylation, we depleted each of the candidate fly acetylases individually in S2 cells using RNAi and assessed levels of acetylated α-tubulin (acTb) by immunoblot (Figure 1 A). Depletion of only one candidate, *CG3967,* resulted in loss of acTb. Thus, we conclude that CG3967 is the major α-tubulin acetylase in S2 cells and refer to it hereafter as dTAT *(Drosophila* α-tubulin acetylase).

**Figure 1.**
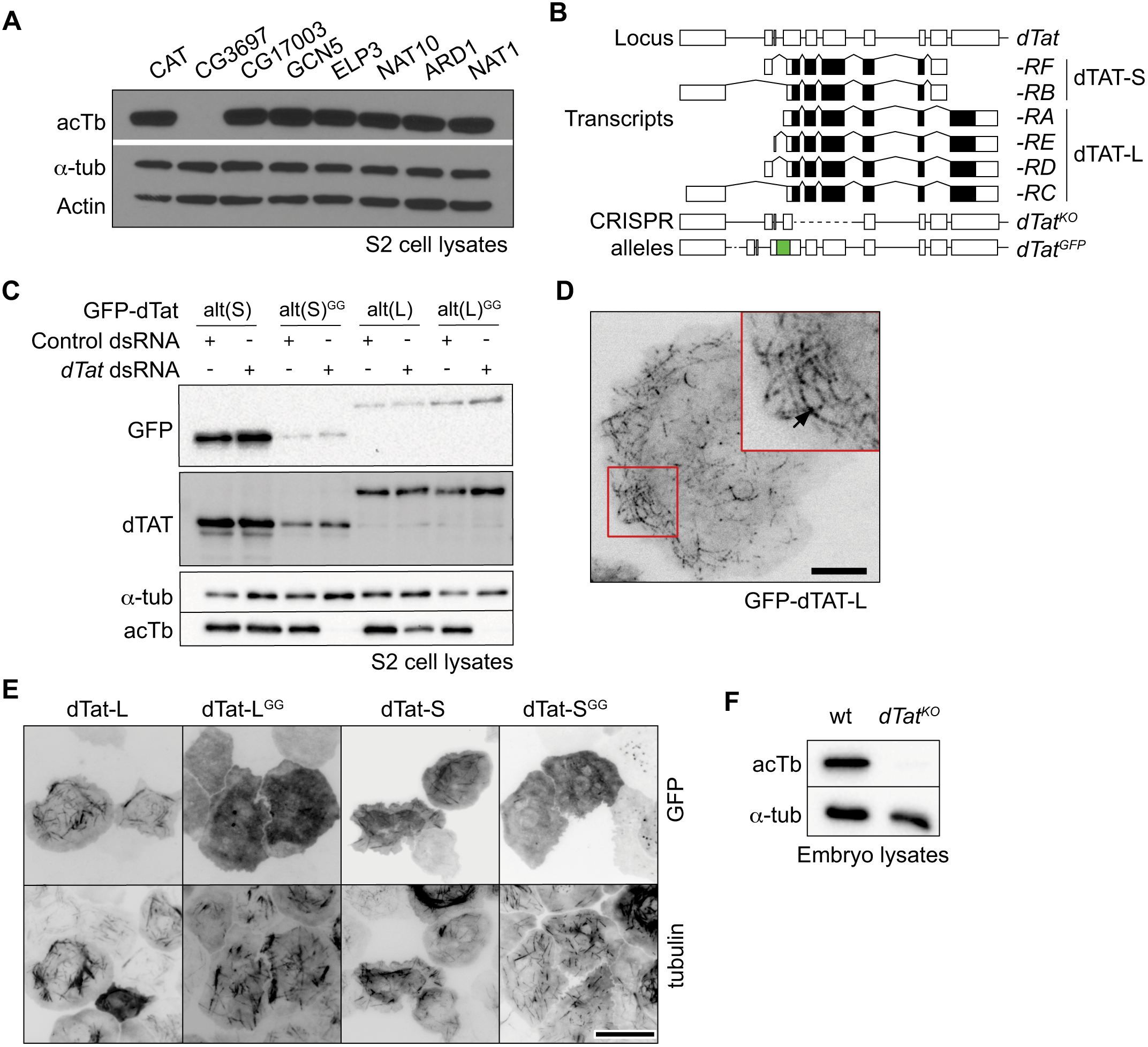
dTAT is the major microtubule acetylase in *Drosophila.* (A) Western blots of cell lysates from S2 cells treated with dsRNA to the indicated genes (CAT, chloramphenicol acetyltransferase dsRNA used as a negative control) showing that *CG3697/dTat* RNAi eliminates acTb immunoreactivity without affecting levels of α-tubulin. (B) Schematic of the *dTat* locus, *dTat* transcripts, and new alleles generated by CRISPR-based genome engineering. Lines depict introns and boxes depict exons (empty boxes, non-coding; shaded, protein coding). The *dTat* locus encodes six documented transcripts encoding two polypeptides (dTAT-L and dTAT-S) with alternative 3’ coding exons. The *dTat*_*KO*_ allele deletes the first three coding exons shared by all *dTat* transcripts and the *dTat^GFP^* allele contains GFP coding sequences fused in-frame upstream of the start codon shared by all isoforms. (C-D) Fluorescence microscopy of S2 cells expressing GFP-dTAT fusions showing the punctate, filamentous localization of GFP-dTAT-L (C) and co-localization of different versions of GFP-dTAT with α-tubulin (D). Scale bar = 5 μm in (C), 15 μm in (D). (E) Catalytic activity of dTAT is required for a-tubulin acetylation. RNAi-resistant versions of GFP-dTAT (GFP-dTATalt) were assayed for their ability to rescue α-tubulin acetylation in *dTat(RNAi)-treated* S2 cells. (F) Western blotting of total protein lysates from wild type and *dTat*_*KO*_ mutant embryos demonstrates that dTAT is required for a-tubulin acetylation *in vivo.*

We used the same strategy to identify the α-tubulin deacetylase. In mammalian cells, both HDAC6 and Sirt2 deacetylate microtubules (Hubbert et al., 2002; North et al., 2003). *Drosophila* has one homologue of *HDAC6* and two related genes, *Sirt1* and *Sirt2,* which exhibit high identity with mammalian *Sirt2.* We targeted HDAC6 and Sirt2 with RNAi and found that acTb levels were only elevated by *HDAC6* dsRNA (Figure S1A). We corroborated these results by immunoblotting samples from *HDAC6*^*KO*^, *Sirt1*^*KO*^, and *Sirt2*^*KO*^ null mutants (Figure S1B). Therefore, our data indicate that α-tubulin acetylation is regulated in *Drosophila* by the antagonistic activities of dTAT and HDAC6 in S2 cells.

The *dTat* coding sequence is predicted to generate two isoforms by alternative splicing: a long (L) isoform encoding a 461 residue (50 kD) protein and a short (S) isoform encoding a 291 residue (30kD) protein (Figure 1B). Both isoforms share an N-terminal 201 residue catalytic domain, but diverge in their C-terminal tails, which lack predicted secondary structure. To define the roles of the two isoforms, we synthesized Myc-tagged RNAi-resistant versions of each isoform using alternate codon usage (*dTat^alt^-L* and *-S*) to make them resistant to RNAi. Both constructs rescued microtubule acetylation in cells treated with dsRNA to deplete endogenous dTAT (Figure 1C). As a control, we made point mutations in both *dTat^alt^* constructs to abolish catalytic activity (G133W and G135W) and found that both inactive mutants failed to rescue microtubule acetylation.

Prior studies have shown that αTAT interacts with microtubules *in vitro* (Akella et al., 2010; Coombes et al., 2016; Kalebic et al., 2013b; Shida et al., 2010; Szyk et al., 2014). Our efforts to localize endogenous dTAT in S2 cells by immunofluorescence with antibodies raised against the catalytic domain were inconclusive. The antibodies labeled punctae that occasionally co-localized with microtubules, but also prominently labeled the cytoplasm as might be expected for a soluble enzyme (data not shown). However, GFP-tagged versions of dTAT^alt^-L and dTAT^alt^-S exhibited robust co-localization with microtubules (Figure 1D, 1E). In contrast, catalytically inactive forms of the enzyme (Topalidou et al., 2012) dTAT^alt^-L^GG^ and GFP-dTAT^alt^-S^GG^ mostly localized in the cytoplasm, suggesting that acetylase activity is required for microtubule interaction.

To study *dTat* function in flies, we used CRISPR/CAS9 to delete the first four exons of the gene to produce a null mutant *(dTat^KO^)* (Figure 1B). Immunoblotting confirmed that *dTat^KO^* homozygous mutants lacked detectable acTb (Figure 1F). We conclude that dTAT is the major α-tubulin acetylase in flies, as in S2 cells.

### dTAT is highly expressed in neurons of the PNS

To define biological functions of dTAT, we first examined expression patterns of *dTat* transcripts. To this end, we isolated different GFP-labeled cell types with FACS and subjected the cell lysates to RNA-Seq analysis (Figure 2A). Among the larval cell types we surveyed, including muscle, epithelia, glia, and neurons, *dTat* expression is highest in neurons. Peripheral nervous system (PNS) neurons, particularly dendrite arborization (da) neurons that mediate responses to mechanical, thermal, light, and proprioceptive stimuli highly express isoforms for both dTAT-S and ‐L, suggesting the PNS is likely an important functional site for dTAT.

**Figure 2.**
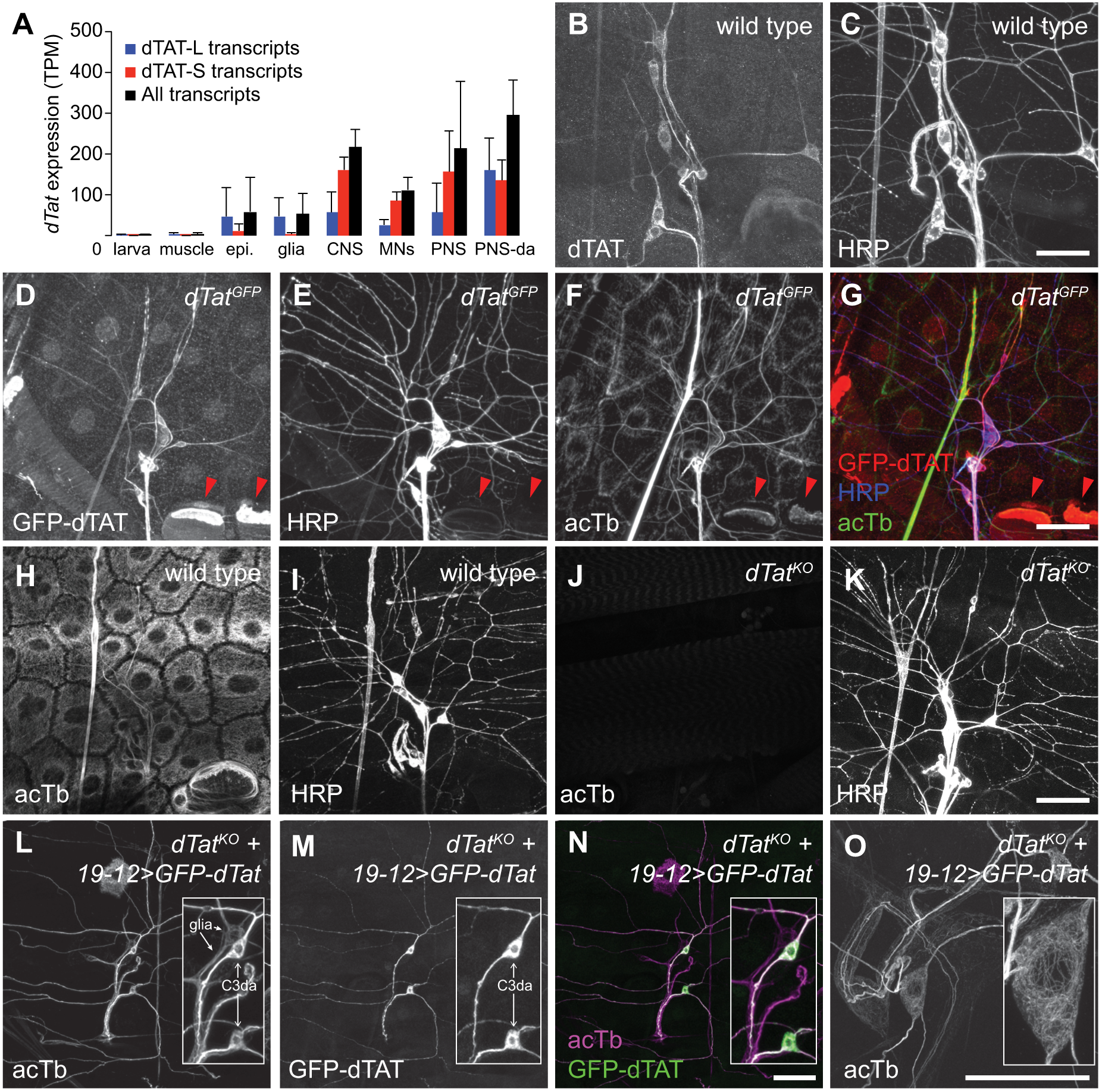
dTAT is enriched in the peripheral nervous system. (A) RNA-Seq analysis of *dTat* expression. GFP-labeled cells of the indicated types were manually picked from enzymatically/mechanically dissociated cell suspensions and subjected to RNAseq analysis. Drivers used to label cells are provided in Table S2. Bars depict mean expression levels of the six predicted *dTat* transcripts in the indicated cell types. tpm = transcripts per million. Error bars = standard deviation. N = 4+ independent samples for each condition. (B-G) Distribution of endogenous dTAT. Maximum intensity projections of larval body walls immunostained with anti-dTAT antibodies (B) revealing that dTAT is enriched larval body wall sensory neurons, which are labeled with antibodies to HRP (C). (D-G) Maximum intensity projections of larval body walls stained with antibodies to (D) GFP to detect dTAT-GFP expressed from a *dTat^GFP^* knock-in allele, (E) HRP to detect sensory neurons, and (F) acTb. dTAT-GFP is enriched in sensory neurons, which exhibit high levels of acTb. (G) Overlay of dTAT-GFP, acTb, and HRP signals. We also note high levels of dTAT-GFP and acTb in apodemes (red arrowheads), consistent with a prior report characterizing acTb distribution (Jenkins et al., 2017). (H-K) *dTat* is required for acTb accumulation *in vivo.* Maximum intensity projections are shown for wild type or *dTat*_*KO*_ mutant third instar larvae stained with antibodies to acTb and HRP. Whereas acTb immunoreactivity is present in sensory neurons and all other body wall cells of control larvae (H, I), acTb signal is absent in *dTat*_*KO*_ mutants (J, K). (L-O) acTb distribution in sensory neurons. Maximum intensity projections are shown for *dTat*_*KO*_ mutant third instar larvae expressing *UAS-GFP-dTat-L* via *19-12-Gal4* and stained with antibodies to acTb (L, O) and GFP (M). Both dTAT and acTb are present throughout the cell body and dendritic arbor of c3da neurons (L, M). (N) Overlay of images from (L) and (M). Insets contain zoomed images of cell bodies and arrows mark the cell types (c3da neurons and peripheral glia) labeled by *19-12-Gal4.* (O) Maximum intensity projection of expanded body wall tissue from *dTat^KO^* mutant third instar larvae expressing *UAS-GFP-dTat-L* via *19-12-Gal4* stained with acTb antibodies. acTb staining forms a continuous network in the soma and dendrites. Scale bars = 50 μm in all panels except (O) where the scale bar = 20 μm in pre-expansion dimensions.

To extend these results, we examined dTAT protein distribution *in situ* using anti-dTAT antibodies. Consistent with our RNAseq results, dTAT immunoreactivity was concentrated in neurons, particularly in PNS-da neurons (Figure 2B-2C). dTAT exhibited elevated accumulation in the soma, axon, and primary dendrites, and undetectable levels in terminal dendrites of da neurons. Likewise, when we monitored GFP-dTAT distribution in larvae homozygous for a GFP knock-in allele (*dTat*^*GFP*^) that produces a GFP-dTAT fusion protein expressed from the endogenous *dTat* locus that supports microtubule acetylation *in vivo,* we found that GFP-dTAT was similarly expressed most highly in PNS neurons (Figure 2D-2G). Within neurons, GFP-dTAT was enriched in the soma, axons, and primary dendrites but largely absent from terminal dendrites.

Given that dTAT is enriched in PNS neurons, we next investigated patterns of acTb accumulation in wild type and *dTat^KO^* mutant larvae. Immunostaining with a monoclonal antibody to acTb revealed a dense network of acTb immunoreactivity in all cells of the body wall, including prominent labeling of PNS neurons (Figure 2H-2I). In contrast, *dTat^KO^* mutant larvae lacked detectable acTb immunoreactivity, demonstrating that *dTat* is required for tubulin acetylation (Figure 2J-2K). Resupplying *dTat* to sensory neurons via Gal4-mediated expression of the *dTat* long isoform containing an N-terminal GFP tag (*UAS-GFP-dTat-L*) rescued microtubule acetylation in a cell-autonomous manner, demonstrating that *dTat* expression in the PNS is sufficient for tubulin acetylation (Figure 2L-2N). This rescue assay additionally allowed us to monitor acTb distribution in PNS neurons as acTb is present only in *dTat*-expressing cells. Similar to GFP-dTAT-L, acTb was enriched in the soma, axons, and primary dendrites of c3da neurons but largely absent from terminal dendrites. Expansion microscopy revealed a dense network of acTb in the soma that extended into axons and dendrites (Figure 2O). Thus, *dTat* is both necessary and sufficient for microtubule acetylation, and acTb distribution mirrors dTAT distribution in neurons.

### dTAT is required for larval mechanosensitivity

Similar to vertebrates, *Drosophila* exhibit several forms of mechanosensation including gentle touch sensation, gravity sensing, mechanical nociception, proprioception and hearing (Gong et al., 2004; Hughes and Thomas, 2007; Kamikouchi et al., 2009; Kernan et al., 1994; Song et al., 2007; Zhong et al., 2010). The microtubule cytoskeleton is involved in mechanosensation, particularly touch responses (Bounoutas et al., 2009; Tanner et al., 1998; Zhang et al., 2015), and mutations in mouse *αTAT1* and *C. elegans mec-17* and *atat-2* reduce mechanosensitivity (Morley et al., 2016; Shida et al., 2010; Topalidou et al., 2012). Furthermore, *Drosophila* gentle touch responses require the function of NOMPC, a TRP channel that relies on microtubule interactions for gating (Zhang et al., 2015). We therefore tested *dTat* mutants for touch sensitivity defects. Gentle touch activates class III da (c3da) neurons to elicit stereotyped behaviors including backward locomotion and turning that can be quantified using a simple and robust scoring matrix (Kernan et al., 1994). As shown in Figure 3A, *dTat* mutation leads to a ∼67% decrease in gentle touch responses. This phenotype was similar in *dTat^KO^* homozygotes and in combination with a deficiency chromosome that spanned the *dTat* locus *(dTat^KO^/Df)* (Figure 3A), suggesting that loss of *dTat* was the root cause of the defects. To test whether this defect in gentle touch responses reflected a requirement for microtubule acetylation, we assayed gentle touch responses of mutants in which lysine 40 (K40) of the major *α-tubulin* isotype *αTub84B,* which accounts for >90% of α-tubulin expression in da neurons (Figure S2), is mutated to a non-acetylatable residue (Jenkins et al., 2017). We found that *αTub84B^K40A^* and *αTub84B^K40R^* mutant larvae exhibited defects in gentle touch responses that were comparable to *dTat^KO^* mutants, strongly suggesting that microtubule acetylation plays a key role in regulating gentle touch responses (Figure 3A). Finally, *dTat^KO^*, *αTub84B^K40R^* double mutant larvae exhibited gentle touch defects that were comparable to either single mutant alone, suggesting that *dTat* and *αTub84B* function in the same pathway for gentle touch responses.

**Figure 3.**
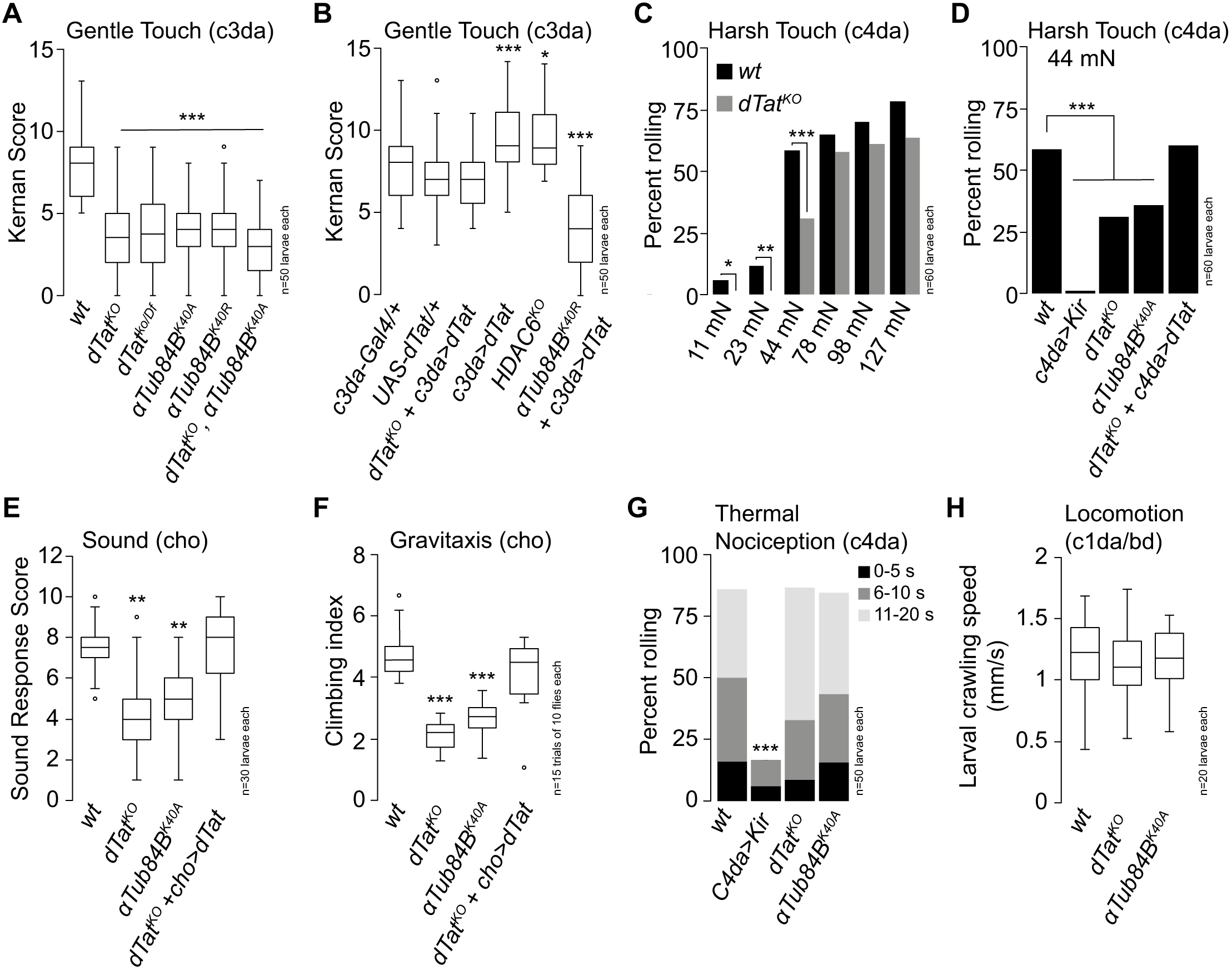
*dTat* regulates *Drosophila* mechanosensation. (A-B) *dTat* is necessary for gentle touch responses. (A) Mutations in *dTat* or *αTub84B* significantly reduce gentle touch responses compared to wild type (wt) controls. Boxplots depict behavioral responses of third instar larvae to gentle touch stimulus at 96 h after egg laying (AEL). In this and subsequent panels, boxes mark 1st and 3rd quartiles, bands mark medians, whiskers mark 1.5 x IQR, and outliers are shown as points. (B) *dTat* expression in c3da neurons rescues larval gentle touch defects and *dTat* overexpression in a wild type background or *HDAC6* mutation enhance touch responses. *P < 0.05, ***P < 0.001 compared to wt controls, Kruskal-Wallis rank sum test with Dunn’s post-hoc test and Bonferroni correction for multiple comparisons. (C-D) *dTat* is necessary for harsh touch responses. (C) Bars depict the proportion of wild type or *dTat* mutant larvae responding to stimulation with von Frey fibers that deliver the indicated amount of force. *dTat* mutants exhibit significant defects in response to 11 mN, 22 mN, and 44 mN stimulus. *P < 0.05, **P < 0.01, ***P < 0.001, compared to wt controls, unpaired t-test with Welch’s correction. (D) *UAS-K’e*r*2*.1 expression in c4da neurons blocks nociceptive responses to 44 mN stimulus, demonstrating that the response is mediated by c4da neurons. *dTat*_*KO*_ or *αTub84B^K40A^* mutation suppress mechanonociceptive responses and *dTat^KO^* mechanonociceptive defects can be rescued by c4da-specific expression of *UAS-GFP-dTat-L.* ***P < 0.001 compared to wt controls, Chi square test. (E-F) *dTat* is necessary for larval sound stimulus responses (E) and adult gravitaxis (F). *P < 0.05, ***P < 0.001 compared to wt controls, Kruskal-Wallis rank sum test with Dunn’s post-hoc test and Bonferroni correction for multiple comparisons. (G) *dTat* is dispensable for noxious heat responses. Bars depict the proportion of larvae that exhibited nociceptive rolling responses to stimulus with a 39.5 °C thermal probe. Responding larvae were grouped in three bins according to response latency: 0-5 sec (black), 6-10 sec (gray), 11-20 sec (light gray). Hyperpolarization of c4da neurons via expression of *UAS-Kir2.1 (ppk>Kir)* blocks nociceptive behavioral responses to noxious heat but mutation of *dTat* or *αTub84B* does not. ***P < 0.001 compared to wt controls, Chi square test. (H) *dTat* is dispensable for larval locomotion. Larvae homozygous for mutations in *dTat* or *αTub84B* exhibit comparable rates of locomotion to wt controls (Kruskal-Wallis rank sum test). The number of larvae tested is shown for each condition.

To determine the site of action for *dTat* in control of gentle touch responses we next performed genetic rescue assays. Expressing *UAS-GFP-dTat-L* in c3da neurons rescued the *dTat* mutant gentle touch defects, demonstrating that *dTat* function in c3da neurons is sufficient to support gentle touch responses (Figure 3B). We also found that overexpressing *UAS-GFP-dTat-L* in c3da neurons of wild type larvae, but not *αTub84B^K40A^* mutant larvae, significantly enhanced gentle touch responses, suggesting that increasing acTb levels potentiates mechanosensitivity in c3da neurons. Consistent with this notion, *HDAC6^KO^* mutant larvae also exhibited heightened gentle touch responses (Figure 3B).

We next asked whether *dTat* was involved in other forms of mechanosensation, including harsh touch, sound stimulus response, and gravitaxis. Harsh touch activates c4da nociceptive neurons to elicit stereotyped nocifensive rolling responses (Zhong et al., 2010), so we stimulated larvae with von Frey filaments and monitored touch-evoked rolling responses as a measure for *dTat* function in harsh touch. *dTat^KO^* mutant larvae exhibited significantly reduced nocifensive responses to stimuli ranging from 11mN-44mN but not 78nM-127mN (Figure 3C), revealing that harsh stimuli can bypass the requirement for acetylation in mechanosensory responses. These mechanonociceptive defects likely reflect a cell-autonomous role for acetylation in c4da neurons as *dTat^KO^* mechanonociception defects were phenocopied by non-acetylatable *αTub84B^K40A^* mutants and rescued by c4da-specific expression of *GFP-dTat-L* (Figure 3D).

Whereas larval responses to gentle and harsh touch are mediated by da neurons, sound and gravity responses are mediated by chordotonal (cho) neurons (Kamikouchi et al., 2009; Zhang et al., 2013). To determine whether *dTat* was also required for mechanosensory responses in cho neurons we next assayed for functions of *dTat* in larval sound stimulus responses and adult gravitaxis. In response to sound stimulus (70dB 500Hz tone), wild type larvae exhibited a stereotyped startle response that was compromised in *dTat^KO^* and non-acetylatable *αTub84B^K40A^* mutants (Figure 3E). As with touch responses, these defects likely reflect cell-autonomous function of *dTat* as expressing *UAS-GFP-dTat-L* in cho neurons of *dTat^KO^* mutants rescued sound startle defects. Finally, adult gravitaxis behavior was impaired in *dTat^KO^* mutants and non-acetylatable *αTub84B^K40A^* mutants and gravitaxis defects of *dTat^KO^* mutants could be rescued by neuronal expression of *UAS-GFP-dTat-L* (Figure 3F). Altogether, these results demonstrate that *dTat* and microtubule acetylation are broadly required for mechanosensation in *Drosophila*.

These mechanosensory defects could reflect a general defect in sensory transduction or a more specific effect on mechanosensation. To differentiate between these possibilities we next assayed for larval responses to noxious heat, which are mediated by c4da neurons – the very same neurons that mediate response to harsh touch (Hwang et al., 2007). Whereas *dTat* mutation compromised harsh touch responses, *dTat^KO^* mutants and non-acetylatable *αTub84B^K40A^* mutants exhibited no marked effect on noxious heat responses (Figure 3G). Furthermore, application of the TRPA1 channel agonist AITC, which directly activates c4da neurons to elicit nocifensive escape responses (Kaneko et al., 2017), generated comparable behavioral responses in wild type controls, *dTat^KO^* mutants and non-acetylatable *αTub84B^K40A^* mutants (data not shown). Finally, we found that wild type controls, *dTat^KO^* and non-acetylatable *αTub84B^K40A^* mutants exhibited comparable rates of larval locomotion (Figure 3H). Taken together, these results demonstrate that *dTat* and microtubule acetylation do not broadly regulate sensory transduction but instead preferentially affect mechanosensory responses.

### dTAT is dispensable for dendrite morphogenesis

How might dTAT influence mechanotransduction? Knockdown of the ARD1-NAT acetylase, which reduces microtubule acetylation, and pharmacological inhibition of the microtubule deacetylase HDAC6, which increases acetylation, compromise dendrite growth in hippocampal cultures *in vitro* (Ageta-Ishihara et al., 2013; Ohkawa et al., 2008). Likewise, mutation of α-Tub84B K40 affects dendrite branching in *Drosophila* c4da neurons (Jenkins et al., 2017). While each of these studies suggest a role for acetylation in dendrite development, both ARD1-NAT and HDAC6 target additional substrates, and α-tubulin K40 may have a structural role and may be subject to additional modifications including methylation (Park et al., 2016). Thus the role of dTAT and α-tubulin acetylation in dendrite development is still unclear.

We therefore explored the possibility that defects in mechanosensory neuron morphogenesis caused by *dTat* mutation might contribute to mechanosensory defects, initially focusing our analysis on NOMPC-expressing mechanosensory c3da and cho neurons. Using *nompC-Gal4* (Petersen and Stowers, 2011) to visualize c3da neurons, which mediate larval responses to gentle touch, we found that *dTat* mutation had no obvious effect on dendrite arborization (Figure 4A-4D) or axon patterning (data not shown). Similarly, when we used antibody staining to label cho neurons, which mediate larval response to sound, we found that *dTat* mutation had no obvious effect on cho neuron morphogenesis (Figure 4E-4G). Thus, *dTat* is dispensable for morphogenesis of NOMPC-expressing larval mechanosensory neurons.

**Figure 4.**
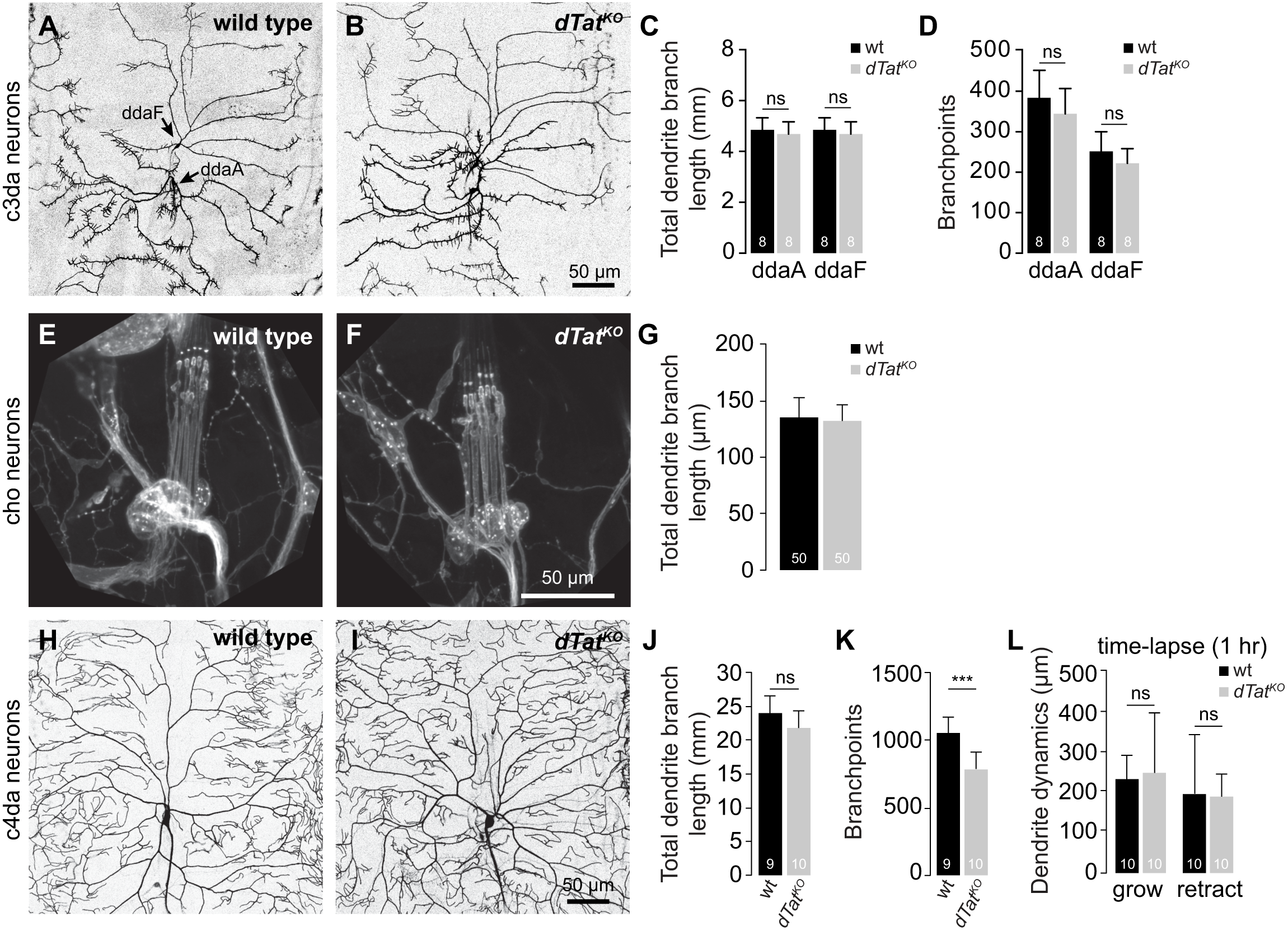
*dTat* is largely dispensable for dendrite morphogenesis in mechanosensory neurons. Representative images of (A) wild type control and (B) *dTat^KO^* mutant c3da neurons labeled with *nompC-Gal4, UAS-mCD4-tdGFP.* Mean and standard deviation are shown for (C) dendrite branch length and (D) number of branch points of dorsal c3da neurons ddaA and ddaF. Representative images of cho neurons from (E) wild type controls and (F) *dTat^KO^* mutant third instar larvae labeled with anti-HRP antibodies. (G) Bar graphs depict mean and standard deviation for cho dendrite length of the indicated genotypes. Representative images of c4da neurons labeled with *ppk-CD8-GFP* are shown for (H) wild type controls and (I) *dTat*^*KO*^ mutant third instar larvae. Mean and standard deviation are shown for (J) dendrite branch length, (K) number of branch points, and (L) the extent of dendrite dynamics measured over a 1 h time-lapse (from 96 h to 97 h AEL). ***P < 0.001, ns, not significant compared to wild type controls, unpaired t-test with Welch’s correction. The number of neurons analyzed for each sample is indicated. Scale bars = 50 μm.

Among *Drosophila* mechanosensory neurons, c4da neurons have the most expansive, highly branched arbors, so we next monitored effects of *dTat* mutation on c4da dendrite growth using the c4da-specific marker *ppk-CD4-tdGFP* (Han et al., 2011) to visualize c4da dendrite arbors in third instar larvae. C4da dendrites establish complete, non-redundant coverage of the body wall (tiling) post-embryonically and maintain this tiling throughout larval development, even as larvae grow (Grueber et al., 2002; Parrish et al., 2009). C4da neurons of *dTat* mutants had no overt defects in the establishment or maintenance of dendritic coverage (Figure 4H, I), in overall dendrite growth (Figure 4J), dendrite growth dynamics (Figure 4L), or axon development (data not shown). However, we did note a minor but consistent reduction in the number of dendritic branch points (Figure 4K), suggesting that *dTat* likely contributes to c4da dendrite development. Prior studies have suggested that microtubule acetylation regulates susceptibility of dendritic microtubules to katanin severing (Sudo and Baas, 2010), therefore we examined the possibility that these minor branching defects might reflect a role for dTAT in regulating katanin-based severing of dendritic microtubules. Using a *UAS-katanin* overexpression assay we found that *dTat* mutation no effect on *katanin*-induced dendrite arbor remodeling (Figure S3). Altogether, these results demonstrate that *dTat* is largely dispensable for mechanosensory neuron morphogenesis, similar to what has been observed in mouse *Atat1* mutants (Morley et al., 2016) and *C.elegans mec-17* mutants (Akella et al., 2010; Shida et al., 2010), suggesting that the wide-ranging effects of dTAT on mechanosensation are likely caused by other mechanisms.

### *dTat* is required for NOMPC-dependent mechanotransduction

The mechanosensory phenotype of *dTat* mutants could reflect defects in mechanosensation or in transmission downstream of mechanosensation. To differentiate between these possibilities, we used Ca^2+^ imaging to measure mechanosensory responses of c3da neurons in wild type and *dTat* mutant larvae. We immobilized semi-intact larval preparations expressing the transgenic calcium sensor GCaMP6s in c3da neurons, provided mechanical stimulus via focal displacement of the larval body wall, and monitored calcium responses in c3da neurons using a confocal microscope (Figure 5A). Consistent with previous reports (Yan et al., 2013), we found that touch stimulus induces robust calcium responses in c3da neurons (Figure 5B). In wild type larvae we observed a progressive increase in calcium responses with increased stimulus strength over a low range of stimuli (10-40 μm displacement), beyond which responses reached plateau (Figure 5C). In contrast, *dTat^KO^* and *αTub84B^K40A^* mutants exhibited significant reductions in touch-induced calcium transients that were most pronounced in the low force range. For example, 30 μm displacement yielded half maximal responses in wild type larvae and negligible calcium responses in *dTat^KO^* and *αTub84B^K40A^* mutants. These defects in calcium responses were progressively attenuated with increased stimulus strength, suggesting that the mechanosensory requirement for *dTat* in c3da neurons can be overcome by increasing stimulus strength, similar to what we observed in c4da neurons (Figure 3C, 5C). These results further suggest that *dTat* functions in PNS neurons to regulate mechanosensory responses.

**Figure 5.**
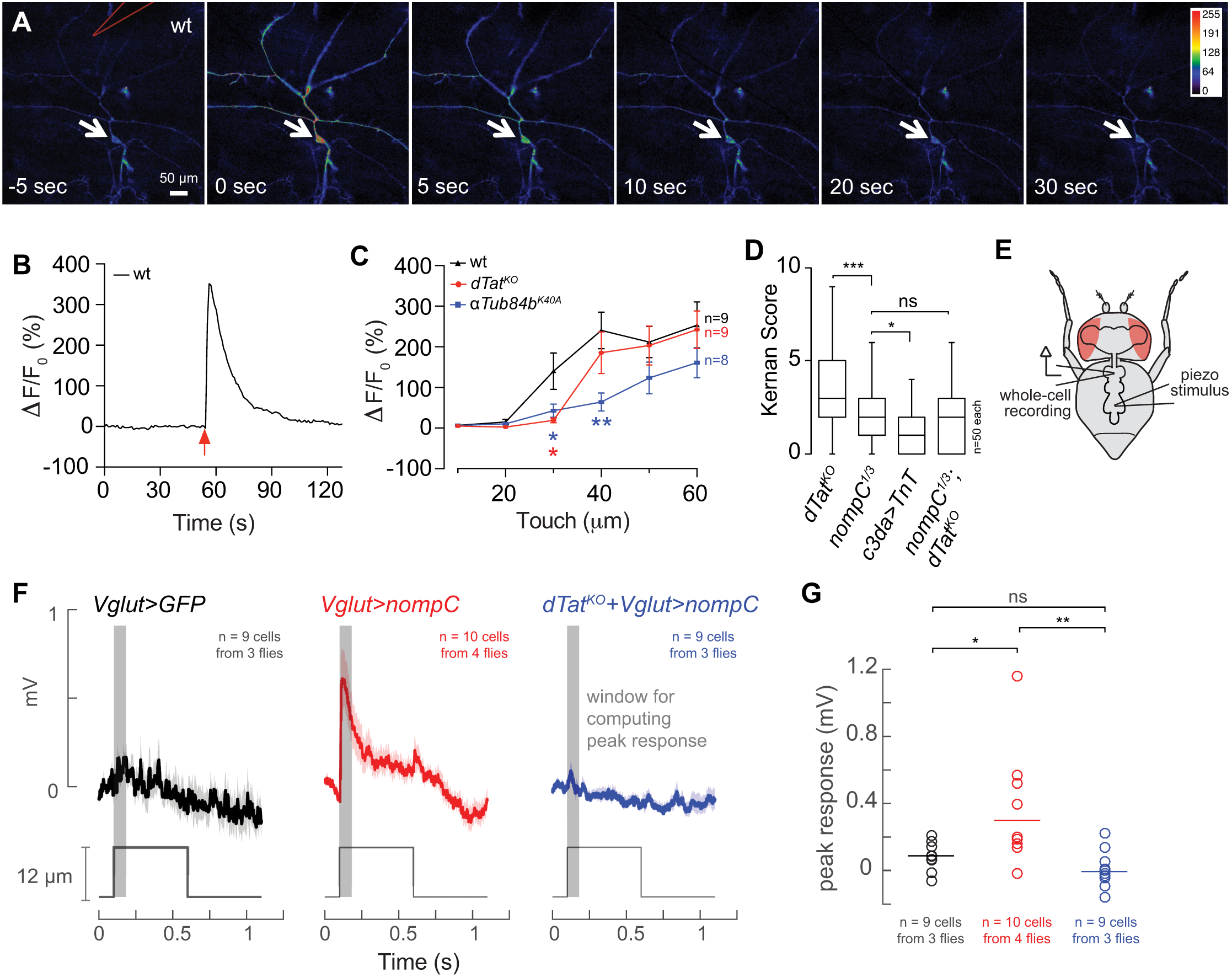
*dTat* is required for NOMPC-mediated mechanotransduction. (A-C) Calcium imaging of c3da neurons responding to mechanical stimulus. (A) Calcium responses to mechanical stimulus delivered via a polished glass electrode (position marked in red, *top left* image) in larvae expressing *UAS-GCaMP6s* in c3da neurons *(nompC-Gal4).* (B) Representative trace of Calcium response measured from the soma of c3da neuron. (C) Mean and standard deviation for Calcium responses [ΔF/F_0_=(F_peak_-F_0_)/F_0_] of larvae of the indicated genotype in response to different mechanical stimuli. *P < 0.05, **P < 0.01, two-way ANOVA followed by Bonfferoni’s multiple comparisons test. (D) *dTat* and *nompC* function in the same genetic pathway for gentle touch. Behavioral responses of third instar larvae to gentle touch stimulus at 96 h AEL are shown for the indicated genotypes. Bar graphs show mean and standard deviation for Kernan scores, and the number of larvae analyzed is shown for each genotype. ***P < 0.001, *P < 0.05, ns, not significant, Kruskal-Wallis rank sum test for multiple independent samples with Dunn’s post-hoc test and Bonferroni correction for multiple comparisons. Brackets indicate pairs being compared. (E-G) *dTat* is necessary for NOMPC-mediated mechanosensory responses. (E) Schematic for recording preparation used in (F, G). (F) Average traces (+/− SEM) from whole-cell patch-clamp recordings for wild type motor neuron expressing *UAS-GFP* (*Vglut>GFP*, *left*), wild type motor neurons expressing *UAS-GFP-nompC* (*Vglut>nompC*, *middle*) and motor neurons from a *dTat^KO^* mutant larva expressing *UAS-GFP-nompC* (*dTat^KO^ + Vglut>nompC*, *right*). Red lines depict mechanical stimulus; note that only the wild type motor neuron expressing *UAS-GFP-nompC* exhibits mechanically-induced depolarization. (G) Scatter plot depicting mean (line) and individual measurements (points) of maximum depolarization minus the baseline resting potential for the indicated genotypes. *P < 0.05, **P < 0.01, ns, not significant, one-way ANOVA followed by Bonfferoni’s multiple comparisons test.

Larval gentle touch responses depend on the TRP channel NOMPC and NOMPC function depends on direct interactions with microtubules, therefore we next conducted genetic epistasis analysis to determine whether dTAT and NOMPC function in the same pathway. While mutation in either *dTat* or *nompC* impaired larval gentle touch responses, *nompC* mutation had more severe effects (Figure 5D). Expressing Tetanus toxin in c3da neurons to block their synaptic output resulted in an even stronger defect than *nompC* mutation (Figure 5D), suggesting that *nompC*-independent pathways contribute to gentle touch responses (Tsubouchi et al., 2012). If *dTat* and *nompC* functioned in independent pathways, we reasoned that the *nompC; dTat* double mutant should have more severe defects than either single mutant alone. Instead, we found that the double mutant was indistinguishable from *nompC* mutants (Figure 5D), suggesting that *dTat* and *nompC* function in the same genetic pathway for gentle touch.

We next tested whether *dTat* was required for NOMPC-mediated mechanotransduction. For these experiments, we first tested whether NOMPC expression could confer mechanosensitivity to neurons that were normally unresponsive to mechanical stimuli. Whereas many PNS neurons including c1da, c3da, c4da, and cho neurons express mechanosensitive ion channels, we found that motor neurons (MNs) do not (Table S1), although MNs do exhibit high levels of *dTat* expression and acTb (Figure 2A, data not shown). Consistent with this expression data, when we conducted *in vivo* whole-cell recordings from leg MNs in the adult ventral nerve cord (Figure 5E), we found that mechanical stimulation of the neuropil did not produce a consistent response (Figure 5F, 5G). By contrast, when we expressed *UAS-nompC-GFP* in leg MNs, mechanical stimulation of the neuropil reliably evoked depolarizing responses. This mechanically-evoked depolarization was abrogated by *dTat* mutation. Thus, NOMPC expression is sufficient to confer mechanosensitivity to MNs and *dTat* is required for NOMPC-dependent mechanotransduction.

We next tested hypotheses for how *dTat* regulates NOMPC function. Microtubule acetylation plays established roles in intracellular transport (Bhuwania et al., 2014; Reed et al., 2006), and NOMPC-microtubule interactions are critical to NOMPC function (Zhang et al., 2015). We thus tested roles for *dTat* in NOMPC localization and in NOMPC-microtubule interactions. We found that *dTat* mutant c3da neurons exhibit normal NOMPC-GFP distribution when NOMPC-GFP is ectopically expressed (Figure 6A). Along with a prior report that ARs are largely dispensable for NOMPC trafficking in c3da neurons (Yan et al., 2013), these findings suggest *dTat* regulates NOMPC function instead of localization.

**Figure 6.**
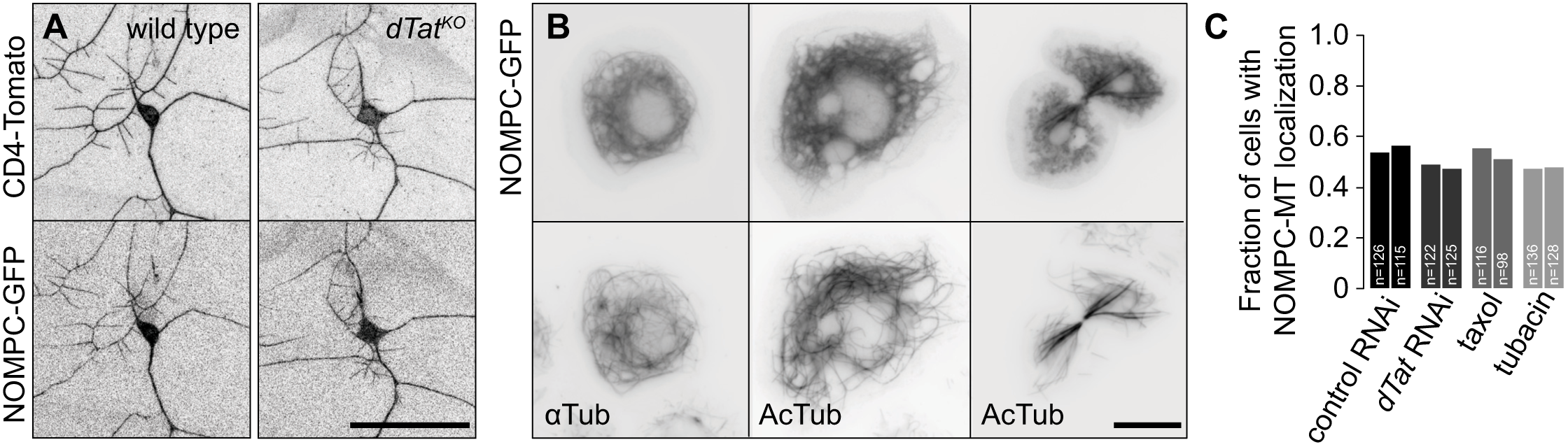
*dTat* is dispensable for NOMPC-microtubule interactions. (A) *dTat* is dispensable for GFP-NOMPC localization. Maximum intensity projections showing GFP-NOMPC distribution in c3da neurons additionally expressing the membrane marker CD4-tdTomato. GFP-NOMPC shows comparable distribution in wild type control (*left*) and *dTat^KO^* mutant (*right*) larvae. (B) NOMPC co-localizes with microtubules in S2 cells. S2 cells stably transfected with *UAS-GFP-nompC* were immunostained with antibodies to GFP and acTb or α-tubulin, as indicated. Images show cells in interphase (*left, middle*) and during anaphase (*right*). (C) NOMPC-microtubule co-localization is unaffected by alteration of acTb levels. S2 cells stably transfected with *UAS-GFP-nompC* were treated with control RNAi, *dTat* RNAi, taxol or tubacin, immunostained using GFP and tubulin antibodies, and the fraction of cells exhibiting NOMPC-microtubule co-localization was visually scored in a blind experiment. Chi-square tests revealed no differences in NOMPC-microtubule co-localization among the different treatments. The number of cells analyzed is shown for each treatment.

When expressed in S2 cells, NOMPC-GFP co-localizes with microtubules and is sufficient to confer mechanosensitivity to this normally non-responsive cell line (Yan et al., 2013). We therefore examined whether NOMPC exhibits a preference for acetylated vs. deacetylated microtubules in S2 cells and found NOMPC-GFP co-localized extensively with acTb (Figure 6B). Although *dTat* RNAi eliminated microtubule acetylation, it did not significantly reduce NOMPC-GFP co-localization with microtubules (Figure 6C). Likewise, treating S2 cells with the HDAC6 inhibitor tubacin or taxol to increase acTb levels had no effect on NOMPC-microtubule co-localization, indicating that acTb is not a critical determinant of NOMPC-microtubule interactions.

### NOMPC-dependent mechanotransduction requires dTAT to prevent breakage of microtubules in C3da neurons

Recent reports have established a role for acTb in controlling mechanical properties of microtubules: loss of acetylation renders microtubules susceptible to mechanical breakage (Portran et al., 2017; Xu et al., 2017). We hypothesized that mechanical stimuli that normally induce bending of microtubules, which then contribute to transduction of mechanical forces, might lead to microtubule breakage in the absence of microtubule acetylation. This model predicts that neuronal microtubules should exhibit increased mechanically-induced breakage in *dTat* mutants and microtubule stabilization should both sensitize wild type larvae to mechanical stimuli and bypass the requirement for *dTat* in mechanosensation.

To test the predictions of this model we first assayed for microtubule breakage using a GFP-tagged version of EB1 *(UAS-EB1-GFP)* to label microtubule plus ends in c3da neurons additionally expressing a membrane-targeted red fluorescent protein *(UAS-CD4-tdTomato).* First, we examined whether *dTat* mutants accumulated microtubule breaks during development. Breaking microtubules, either by severing enzymes or mechanical forces, generates additional microtubule ends, and new growth from the microtubule plus-ends can be monitored using EB1-GFP. Thus, an increase in EB1-GFP puncta correlates with an increase in microtubule breakage. In third instar larvae, *dTat* mutant c3da neurons exhibited ∼70% more EB1-GFP-positive puncta than controls (control, 1.74 ± 0.51; *dTat*, 2.99 ± 0.90), possibly reflecting breakage induced by larval peristalsis, contact with the substrate, or other mechanosensory stimuli that larvae encounter (Figure 7A-7E). Next, we examined whether microtubules in *dTat* mutants were more prone to mechanically-induced breakage by imaging EB1-GFP immediately before and after mechanical stimulus. We found that mechanical stimulus induced a significant increase in EB1-positive puncta in c3da neurons of *dTat* mutants but not wild type controls (Figure 7F-7H).

**Figure 7.**
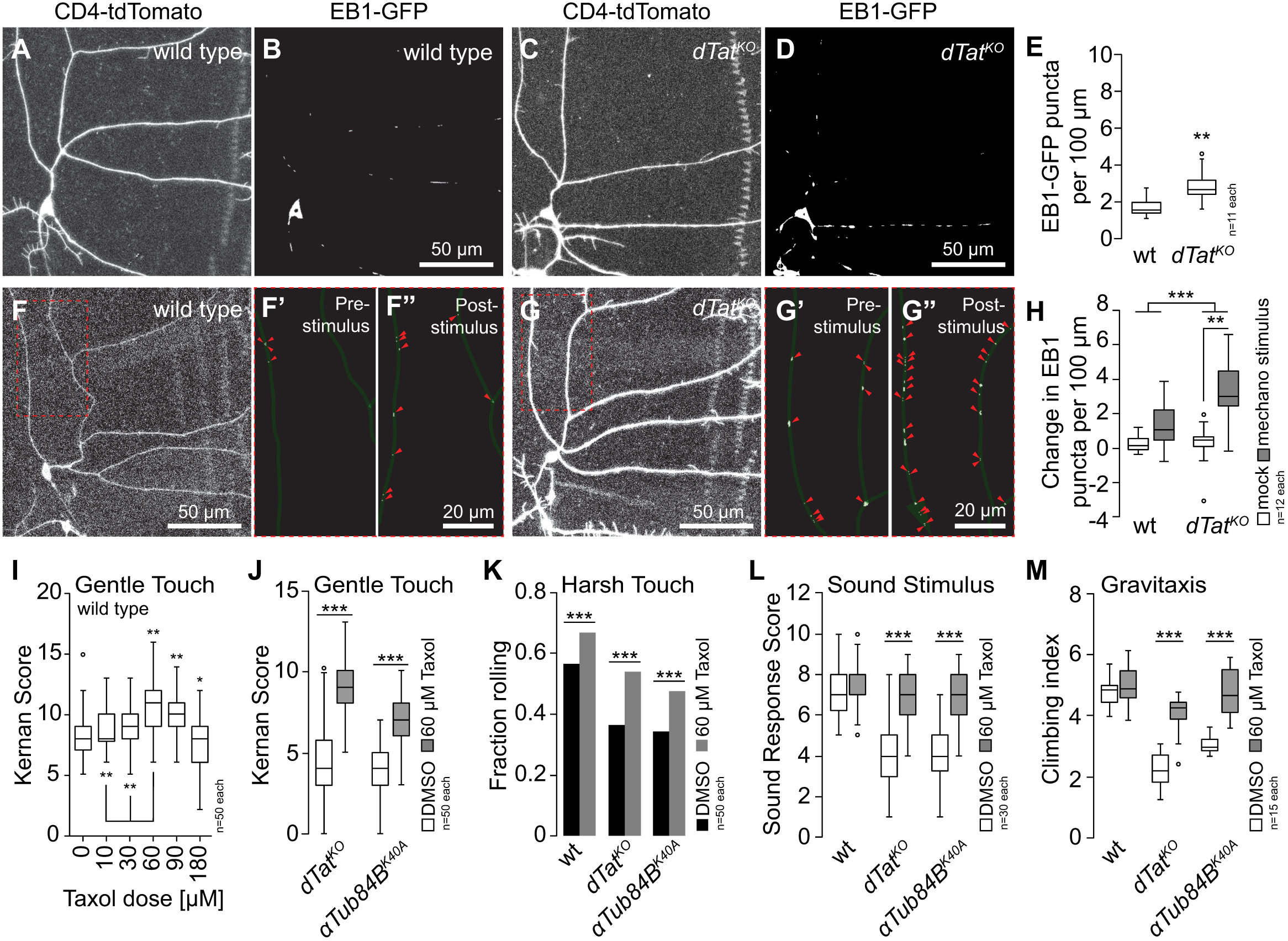
*dTat* promotion of microtubule mechanical stability modulates mechanotransduction. (A-E) *dTat* mutants accumulate microtubule breaks. Representative images of c3da neurons from (A, B) wild type and (C, D) *dTat^KO^* mutant larvae expressing EB1-GFP to label microtubule plus ends and the membrane marker CD4-tdTomato. (E) Box plot depicts the number of EB1-positive puncta per 100 mm of dendrite length. **P < 0.01 compared to wild type control, unpaired t-test with Welch’s correction. (F-H) Microtubules of *dTat^KO^* mutants are prone to mechanically-induced breakage. Images of representative (F) wild type and (G) *dTat* mutant c3da neurons that were subjected to mechanical stimulus are shown. EB1-GFP labeling is shown immediately before (F’ and G’) and after stimulus (F” and G”), and EB1 puncta are marked with red arrowheads. (H) Box plot depicting quantification of EB1-positive puncta. Two-way ANOVA analysis revealed a significant interaction effect between mechanical treatment and genotype on EB1 puncta number F(2,12) = 7.539, P = 0.009. Simple main effects analysis showed a significant difference in EB1 puncta number between treated and untreated *dTat* mutant larvae (P = 0.001) but not between treated and untreated wt larvae (P = 0.184). (I-M) Taxol-induced microtubule stabilization rescues mechanosensory defects of acetylation mutants. (I) Box plot depicting gentle touch response of larvae fed the indicated dose of taxol for 3 h. **P < 0.01, *P < 0.05 compared to vehicle-fed controls, one-way ANOVA with a post-hoc Dunnett’s test. (J-L) Larvae of the indicated genotypes were fed vehicle control or 60 μM taxol for 3 h. (J) Box plot depicting gentle touch response of *dTat* and *αTub84b* mutant larvae fed vehicle control or 60 μM taxol. (K) Bars depict the proportion of larvae of the indicated genotype that exhibited a nociceptive rolling responded to 44 mN von Frey stimulus. (L) Box plot depicting sound stimulus responses of wild type, *dTat* and *αTub84B* mutant larvae. (K) Box plot depicting gravitaxis behavior of wild type, *dTat* and *αTub84B* mutant flies fed vehicle control or 60 μM taxol for 12 h. ***P < 0.001 compared to vehicle controls, unpaired t-test with Welch’s correction for (J-M).

The increased susceptibility of *dTat* mutant microtubules to mechanical breakage likely limits the ability of these microtubules to transduce mechanical forces, thus inhibiting the gating of NOMPC. Our finding that *dTat* overexpression and *HDAC6* mutation enhanced gentle touch responses suggested that stabilizing microtubules in an otherwise wild type background might be sufficient to enhance gentle touch sensitivity. To test this possibility, we examined whether taxol stabilization of microtubules potentiated behavioral responses to gentle touch. Indeed, we found that acute taxol feeding led to dose-dependent increases in gentle-touch sensitivity in third instar larvae over a range of 0-60 μM taxol (Figure 7I) with no obvious effects on neuron morphology (data not shown). Mutation of *nompC* rendered taxol-fed larvae insensitive to gentle touch as did expression of tetanus toxin specifically in c3da neurons (Figure S4), indicating that the increased responses to gentle touch require *nompC* and reflect enhanced activity of c3da neurons.

To test whether mechanosensory defects of *dTat^KO^* mutants and non-acetylatable *αTub84B^K40A^* mutants reflect an increased susceptibility of microtubules to mechanical breakage, we treated larvae with vehicle or 60 μM taxol and measured behavioral responses to gentle touch (Figure 7J). Remarkably, taxol feeding significantly enhanced gentle touch response of *dTat^KO^* and *αTub84B^K40A^* mutants. Likewise, taxol feeding potentiated larval responses of wild type larvae to harsh touch and significantly enhanced harsh touch response of *dTat^KO^* and *αTub84B^K40A^* mutants (Figure 7K). Finally, although taxol feeding did not enhance sound startle responses or gravitaxis in wild type controls, it did restore these behaviors to wild type levels in *dTat^KO^* and *αTub84B^K40A^* mutants (Figure 7L-7M). Altogether, these results support a model in which microtubule acetylation by *dTat* broadly regulates mechanosensation via effects on microtubule mechanical stability.

## Discussion

Microtubule acetylation is required for touch sensation in several model systems (Akella et al., 2010; Morley et al., 2016), however, the molecular mechanisms that underlie its role in mechanosensation are poorly understood. In this study, we identified the major α-tubulin acetylase in *Drosophila,* dTAT, and found that it is broadly required for mechanoreceptivity. In response to gentle touch, *dTat* functions in the same pathway as the TRP channel *nompC* and is required for NOMPC-dependent touch-evoked neuronal responses. Loss of *dTat* causes an increase in microtubule plus ends in the dendrites of somatosensory neurons and taxol stabilization of the microtubule cytoskeleton rescues mechanoreceptivity, suggesting that microtubule breakage underlies the loss of touch sensitivity. Collectively, our results suggest a model in which K40 acetylation functions to stabilize the microtubule cytoskeleton to promote NOMPC activation through its cytoplasmic microtubule-associated tension gate. We speculate that microtubule acetylation likewise promotes activation of other mechanosensory channels by facilitating microtubule-mediated mechanotransduction.

To date, K40 acetylation has been implicated in a variety of cellular processes including cell motility, autophagy, adhesion, intracellular trafficking, and touch sensation (Aguilar et al., 2014; Geeraert et al., 2010; Kalebic et al., 2013a; Montagnac et al., 2013; Reed et al., 2006). In these studies, α-tubulin acetylation was perturbed by expression α-tubulin K40 mutants that could not be acetylated, by manipulating levels of αTAT/MEC-17, or by inhibition of the tubulin deacetylases HDAC6 and Sirt2 using genetic tools and small molecule inhibitors. Interpretation of these results can be complicated by several factors. For example, several groups have described a non-catalytic activity for αTAT/MEC-17 in regulating microtubule dynamics (Kalebic et al., 2013b), making it difficult to parse the role of K40 acetylation from other, poorly understood mechanisms in loss-of-function experiments. In addition, both HDAC6 and Sirt2 deacetylate multiple cytoplasmic substrates, some of which are cytoskeletal proteins, and histones to regulate gene expression through epigenetic mechanisms. Our results indicate that α-tubulin acetylation in *Drosophila* is controlled by the antagonistic activities of dTAT and HDAC6. Although we have not directly examined cell motility, autophagy, or cell adhesion in *dTat* mutants, the fact that they are viable, develop normally, and have normal lifespan suggests that these processes are minimally perturbed in *Drosophila*. However, our results corroborate the role for K40 acetylation in touch perception described in mice and worms (Akella et al., 2010; Morley et al., 2016), indicating that this is a conserved neuronal function. Moreover, our results demonstrating defects in sound perception and gravitaxis in *dTat* mutants reveal that microtubule acetylation plays a broader role in mechanosensation than was previously recognized in other model systems; elucidating the mechanistic basis for this expanded role in nervous system function will be an important goal for future work.

The results of our genetic, electrophysiological, and cell biological experiments indicate that *dTat* is necessary for mechanosensation by the TRP channel NOMPC. NOMPC has 6 central membrane-spanning α-helices and an elongated N-terminal cytoplasmic domain containing 29 tandem ARs. The NOMPC AR domain interacts with the sides of microtubules, forming a link to the NOMPC pore regions and is thought to function as an elastic tether that allows mechanical membrane deformation to gate channel opening in response to mechanical force. Further, *dTat* and *nompC* mutants exhibited similar defects in touch sensitivity when assayed by behavior. Collectively, our data support a model in which dTAT is necessary for touch-evoked responses downstream of NOMPC in c3da and motoneurons. This requirement for *dTat* is not due to NOMPC trafficking and unlikely due to interactions with microtubules, however, we cannot rule out the possibility that other neuron-specific post-translational modifications, neuronal tubulin isoforms, or MAPs could contribute to NOMPC-microtubule interactions.

Intriguingly, loss of *dTat* does not phenocopy *nompC* null mutants in all respects. NOMPC functions in proprioceptive c1da neurons to control larval locomotion (Cheng et al., 2010), but *dTat* mutant larvae exhibit normal locomotion. We speculate that this may be due to neuron class-specific susceptibility to loss of acTb. For example, the microtubule network in c1da neurons might be more resistant to loss of *dTat* than in c4da neurons. We also discovered that adult gravitaxis is perturbed in *dTat* mutants. While NOMPC is dispensable for gravitaxis, TRPA and TRPV family channels are required for gravity-sensing (Kamikouchi et al., 2009; Sun et al., 2009). Likewise, *dTat* mutants have defects in harsh touch responses, which involve TRP, Piezo, and Ppk/ENaC channels but not NOMPC, as well as defects in larval hearing, which involves several TRP channels in addition to NOMPC. These results raise the interesting possibility that interactions between acTb and other channels may play important roles in mechanosensation.

Given the role for *dTat* in NOMPC-dependent touch perception, what is the molecular mechanism? Over the last year, a remarkable series of studies from the Théry and Nachury labs have implicated α-tubulin acetylation as an adaptive biochemical mechanism that allows microtubules to resist mechanical stress (Portran et al., 2017; Xu et al., 2017). These labs developed *in vitro* assays in which individual microtubules could be mechanically stressed by repeated bending and release while imaged by fluorescence microscopy. Cycles of bending induced acute damage to the polymer resulting in decreased microtubule stiffness and localized material fatigue. Using these assays, the authors showed K40 acetylation enhanced microtubule flexibility and increased their mechanical resilience. They found that acetylation weakened inter-protofilament interactions, allowing microtubules to comply with deformative forces without breaking. We favor microtubule breakage as the mechanism underlying loss of mechanosensitivity in *dTat* mutants; we suspect that in the absence of K40 acetylation, microtubules in c3da neurons are mechanically damaged thereby decreasing NOMPC-microtubule interactions and inhibiting the channel’s ability to transduce mechanical stimuli. We speculate that the peristaltic contractility that drives larval locomotion is sufficient to damage microtubules in c3da neurons in *dTat* mutants, whereas in wild type animals acetylated microtubules are able to resist these compressive forces. Consistent with this model, we observed an increase in EB1-labeled microtubule plus ends that could be the result of breaks in *dTat* mutants relative to wild type and showed that stabilizing the cytoskeleton by feeding mutant animals taxol rescued touch sensitivity.

Prior studies into the role of microtubule acetylation in touch perception have arrived at molecular mechanisms that differ from our model, but are not mutually exclusive. In *C. elegans* touch receptor neurons, the sensory dendrite is packed with a cross-linked bundle of long, specialized 15-protofilament microtubules that are specific to this cell type (Chalfie and Sulston, 1981). Mutation of the two α-tubulin acetylases *mec-17* and *atat-2* results in the loss of these unique microtubules and insensitivity to touch (Bounoutas et al., 2009). Thus, K40 acetylation in this system is required for the assembly of a specialized population of microtubules involved in mechanotransduction. Although there is no evidence for specialized microtubules in *Drosophila* da neuron dendrites, the results from worm touch receptors mirror our results in that microtubule acetylation is required to maintain an intact microtubule network. In mouse peripheral sensory neurons, acetylated microtubules are enriched in a sub-cortical band in the soma that was distinct from the cytoplasmic network. Morley et al. showed that this sub-membranous band confers increased rigidity to the plasma membrane and that neurons from *Atat1* knock-out mice exhibited increased membrane stiffness compared to wild type (Morley et al., 2016). The authors proposed that, in this system, microtubule acetylation functions to tune the mechanical properties of the membrane and, in its absence, cells are less elastic and require more force to trigger mechanosensitive channels. Although there is no evidence for a specialized network of acetylated microtubules in *Drosophila* PNS neurons, it will be interesting to determine if *dTat* plays a role in regulating their mechanical properties in the future. It is possible that microtubule acetylation regulates mechanosensitivity through multiple mechanisms by directly regulating microtubule structure and indirectly through tuning mechanical properties of neurons.

## Materials and Methods

### Fly stocks

Flies were maintained on standard cornmeal-molasses-agar media and reared at 25° C under 12 h alternating light-dark cycles. The following fly lines were used in this study: *w*^*1118*^(BL6326); *w*^*1118*^; *ppk-Gal4* (BL32079); *w*^*1118*^, *ppk-mCD8-GFP* (Jiang et al., 2014). *UAS-GFP-dTat-L* (this study), *dTat*^*KO*^ (this study), *dTat*^*GFP*^ (this study), *Df(3L)BSC113* (BL8970), *αTub84B-K40A* (Jenkins et al., 2017), *αTub84B-K40R* (Jenkins et al., 2017), *αTub84B*^*Δ*^ (Jenkins et al., 2017), *hdac6*^*Δ*^, *nompC*^*1*^, *nompC*^*3*^, *Actin-Gal4*, *Gal419-12*, *Gal421-7*, *nompC-gal4*, *ppk-Gal4*, *UAS-TnT*, *UAS-Kir2.1*, *UAS-EB1-GFP*, *UAS-CD4-tomato*, *ppk-CD4-tdTomato*. A list of experimental genotypes is available in Table S2.

### Generation of *dTat* alleles

The *dTat*^*KO*^ and *dTat*^*GFP*^ alleles were engineered using CRISPR/Cas9 mediated gene editing. Target sites were selected upstream of the first coding exon and in the second intron of *dTat* using the CRISPR Optimal Target Finder (http://tools.flycrispr.molbio.wisc.edu/targetFinder). chiRNA plasmids were generated by annealing sense and antisense gRNA oligos, digesting with BbsI, and ligating into the pU6-BbsI-chiRNA expression vector. Donor vectors were generated by cloning homology arms into pHD-DsRed. For *dTat^KO^*, homology arms were designed to delete an 869 base pair fragment spanning the first three coding exons, beginning 16 base pairs upstream of the start codon. The donor vector for *dTat^GFP^* was generated as follows: pHD-dsRed was digested with EcoRI and the 5’ homology arm corresponding to sequences immediately upstreat of the *dTat* start codon, GFP and *dTat* PCR fragments were cloned into the backbone by Gibson assembly (NEB Gibson Assembly Kit) with GFP fused in-frame with the N-terminus of *dTat* and the LoxP-flanked 3xP3-dsRed marker within the second intron. The newly assembled plasmid was then digested with XhoI and the 3’ homology arm was inserted using Gibson assembly. All primer sequences are available in Table S3.

chiRNA and pHD-DsRed plasmids were co-injected into embryos expressing Cas9 in the germline (BL55821: *y[1] M{vas-Cas9.RFP-}ZH-2A w[1118]/FM7a, P{w[+mC]=Tb[1]}FM7-A*), stocks were established from RFP-positive founder males, and 3xP3-RFP markers were removed using Cre-mediated reduction (BL34516: *y[1] w[67c23] P{y[+mDint2]=Crey}1b; sna[Sco]/CyO; Dr[1]/TM3, Sb[1]*) as previously described (Gratz et al., 2014).

### RNA-Seq of larval cells

Four to seven samples of one hundred cells each were isolated and RNA-Seq libraries were prepared as described previously (Boiko et al., 2017). Briefly, third instar larvae expressing *UAS-Red Stinger* in the target cell type (epithelia, *a58-Gal4*; muscles, *mef-Gal4*; peripheral neurons, *elav-Gal4*; central neurons, *elav-Gal4*; motor neurons, *ok371-Gal4*; central glia, *repo-Gal4*) were dissected to isolate the tissue of interest. Bodywalls or brains were dissociated and Red-Stinger-labeled cells were isolated by flow cytometry into RNAqueous lysis buffer. Samples were sequenced as 51 base single end reads on a HiSeq 2500 running in high-output mode at the UCSF Center for Advanced Technology, with read depths ranging from 1.5 to 18.4 million reads. Reads were demultiplexed with CASAVA (Illumina) and read quality was assessed with FastQC (http://www.bioinformatics.babraham.ac.uk/projects/fastqc). Reads were aligned to the *D. melanogaster* transcriptome, FlyBase genome release 6.10, using STAR version 2.5.2b (57) with the option ‘--quantMode TranscriptomeSAM’. Transcript expression was modeled from these STAR alignments using Salmon in alignment-based mode. The raw sequencing reads and gene expression estimates are available in the NCBI Sequence Read Archive (SRA) and in the Gene Expression Omnibus (GEO), accession number pending. The da neuron and whole larvae data sets were previously described and published and are available from SRA and GEO under accession numbers GSE72884 and GSE99711, respectively.

### Plasmid Constructs

Expression constructs for dTAT were generated using PCR from a synthetic template (dTATalt) in which the wobble position of each codon was conservatively replaced, making them refractory to RNAi (Blue Heron Biotech). The GFP-dTATalt fusions were cloned by 5’ overlapping extension PCR using primers listed in Table S3. GFP-dTATalt inserts were cloned into the KpnI/ApaI sites of pMT B V5/His (Invitrogen) for copper-inducible expression in S2 cells or into the EcoRI/NotI sites in pBID-UASC (Addgene) for generating transgenic fly lines. The catalytically inactive version of dTAT was generated by mutating two glycine residues (G133 and G135) to tryptophan to disrupt acetyl CoA binding (Topalidou et al., 2012).

### S2 cell culture, RNAi, and immunofluorescence

Culture and RNAi of *Drosophila* S2 cells were performed as previously described (Rogers and Rogers, 2008). S2 cells (Drosophila Genomics Resource Center, Bloomington, IN) were cultured in SF900II medium supplemented with 100x antibiotic-antimycotic (Invitrogen, Carlsbad, CA). DNA templates for dsRNA synthesis were obtained by PCR amplification of the pFastBacHT-CAT expression plasmid (Invitrogen), BDGP cDNA clones, or S2 cell genomic DNA using the gene-specific primer sequences (Table S3). As a negative control, a sequence from chloramphenicol acetyltransferase (CAT) was amplified and transcribed into dsRNA. In vitro transcription reactions were performed with T7 RNA polymerazse purified in house. Cells were transfected using Fugene HD (Promega) according to the manufacturer’s instructions. Stable cell lines were selected by supplementing culture medium with 10 μg/mL blasticidin or 500 ug/mL hygromycin (Invitrogen). Immunofluorescence was performed by plating cells into fabricated 35 mm glass bottom culture dishes pre-coated with concanavalin A in serum-free Schneider’s medium (Sigma). After cells had attached and spread for 1 hour, they were fixed with 10% formaldehyde (EM Sciences) in HL3 buffer (70 mM sodium chloride; 5 mM potassium chloride; 20 mM magnesium chloride hexahydrate; 10 mM sodium bicarbonate; 5 mM trehalose; 115 mM sucrose; 5 mM HEPES; pH 7.2) for 10 minutes. Cells were permeabilized and blocked with 5% bovine serum albumin in TBST (TBS + 0.1% Triton X-100) before staining with primary and secondary antibodies. Cells were imaged on an Eclipse Ti-E microscope with a 100x oil NA-1.45 objective, driven by NIS Elements software. Images were captured with a cooled charge-coupled device camera (CoolSNAP HQ, Roper Scientific).

### Antibodies

The following antibodies were used in this study: anti-acTb (6-B11-1, Sigma), anti-α-tubulin (DM1α, Sigma), FITC-labeled DM1α monoclonal antibodies (Sigma), anti-GFP (A-11122, Fisher), anti-DsRed (632496, Clontech), Anti-Myc (9E11, DSHB), Anti-Futsch (22C10, DSHB), Cy5-conjugated anti-HRP (Jackson immunoresearch), and Alexa-fluor conjugated secondary antibodies (Fisher). In order to generate antibodies against dTAT, we used PCR to amplify the catalytic domain (residues 1 to 196) and cloned this fragment into the NheI/ XbaI sites of pET28a or the BamHI/NotI sites of pGEX6P2. Recombinant dTAT 1-196 was expressed in *E. coli* and purifiied on NiNTA resin (Qiagen) and glutathione-sepharose, respectively. Purified 6xhis-dTAT1-196 protein was used to generate polyclonal antibodies in rabbits (Pocono Rabbit Farm) and the antibodies were further affinity-purified on GST-dTAT 1-196 bound to amino-link resin (Thermo Fisher Scientific). Secondary antibodies for immunofluorescence were purchased from Jackson Immunoresearch. HRP-conjugated secondary antibodies for immunoblots were purchased from Sigma.

### Imaging of larval samples

#### Live imaging

Larvae were mounted in 90% glycerol under No. 1 coverslips and imaged using a Leica SP5 microscope with a 40× 1.2 NA lens. For time-lapse analysis, larvae were imaged at the indicated time, recovered to yeasted agar plates with vented lids, aged at 25°C, and imaged again.

#### EB1 assays

Larvae were carefully mounted and imaged, then immediately placed in a small plastic petri dish and stimulated with forceps pinches to segments A2, A3, and A4 and re-imaged under the same conditions. Maximum projections of confocal z-stacks (1 μm z step size) were set to identical threshold levels to eliminate non-punctate signals and quantified in ImageJ.

#### Immunostaining

Larval immunostaining was performed as described (Grueber et.al, 2002) with the exception of fixation in freezing methanol for 15 min for AcTb staining. Antibody dilutions were as follows: acTb, 1:1000; GFP, 1:100; 22c10 (1:200); HRP-Cy5 (1:100), secondary antibodies (1:200).

#### Calcium imaging

Third-instar larvae were dissected in calcium imaging solution (310mOsm, pH7.2) containing (in mM): NaCl 120, MgCl_2_ 4, KCl 3, NaHCO3 10, Glucose 10, Sucrose 10, Threalose 10, TES 5, HEPES 10. Larvae were pinned ventral side up on Sylgard^®^ 184 silicone elastomer plates. After opening the larval body from the ventral side, internal organs were removed, and the muscle covering the dorsal c3da neuron (ddaF) was also gently removed to facilitate imaging of ddaF in segments A4 to A6.

Stimulation electrodes (sealed/polished with a diameter of ∼10μm) were mounted in contact with the internal side of the larval body wall within the dendritic filed of the target c3da neuron. The tip position was set to avoid direct contact with dendritic tips for potential damage. Following 20ms transient vertical stimulations, each with increasing displacements (10, 20, 30, 40, 50 and 60μm; applied with a Sutter MP-285 micromanipulator), the electrode was returned to the starting position. GCaMP6s fluorescence was excited with a 488nm solid-state laser and GCaMP fluorescence was imaged at a 0.97Hz frame acquisition rate using a W Plan-APOCHROMAT 20×/1.0 objective lens and a Zeiss LSM 700 confocal microscope. Changes in calcium levels in the cell body were measured using the following formula:

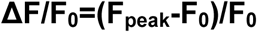

where **F_o_** is the average fluorescence in 30s right before vertical mechanical stimulation. **F_peak_** is defined as the maximum fluorescence upon stimulation.

### Behavior assays

#### Gentle touch

Each larva was stimulated with an eyelash stroke on thoracic segments while in a bout of linear locomotion. The stimulus was applied and scored four times per larva, with responses scored as previously described (Kernan et.al 1994). Tests were performed with the experimenter blind to genotype in this and all other behavior assays.

#### Harsh Touch

Larvae were placed in a plastic petri dish with enough water, so larvae remained moist, but not floating in the dish. Von frey filaments made from fishing line and affixed to glass capillaries were applied to the dorsal side of the larvae between segments A3-A6 until the filament buckled, exhibiting a pre-determined force. Forces of 44mN, 78mN and 98mN were used in this study. A positive response was scored if one complete nocifensive roll occurred after the first mechanical stimulus.

#### Sound/vibration

Wandering third instar larvae were picked from a vial and washed with PBS. 10 larvae were placed on a 1% agar plate on top of a speaker and stimulated as previously described (Zhang et al., 2013). A 1-second 70dB, 500Hz pure tone was played 10 times with 4 seconds of silence in between. Video recordings captured larval behavior, with the number of times out of 10 each larva exhibited sound startle behavior as its individual score. 3 separate trials were performed for each genotype. Videos were recorded with AmScope MU300 Microscope digital camera. Larval startle behavior was scored as responsive with the following behaviors: mouth-hook retraction, pausing, excessive turning, and/or backward locomotion.

#### Gravitaxis

A RING apparatus was assembled as described (Kamikouchi et al., 2009) using 2.3 cm diameter, 9.5 cm polypropylene *Drosophila* vials. 25 flies were collected, aged until all the flies were 5-8 days old, and transferred to a gravitaxis vial sealed with parafilm. The apparatus was rapped on a table five times in rapid succession to initiate the gravitaxis response. Videos of the flies were captured and position of each fly in the tube was determined 4 seconds after the response initiation. Flies were allowed to rest 1 minute, and this was repeated for 5 total trials, for n=1. Assays were repeated for each genotype for total n=3

#### Thermal Nociception

Local heat probe assays were performed as previously described (Chattopadhyay et al., 2012). Washed larvae were placed on a piece of vinyl and stimulated on their dorsal midline at segment A4 with a thermal probe maintained at 38 °C for a maximum of 20s or until completion of a rolling nocifensive response.

#### Larval locomotion

Larvae were washed and placed on agar plate, 10 second videos of individual crawling larvae were recorded in LAS as uncompressed avi files. Files were converted to flymovieformat with any2ufmf and analyzed in Ctrax (Branson et al. 2009). Videos were recorded on Leica DFC310 FX camera on an AmScope FMA050 mount.

#### Taxol Feeding

Third instar larvae were transferred to 35 mM dishes of cornmeal-molasses food containing DMSO (vehicle) or 60uM taxol (unless otherwise indicated) for 3 hours and then subject to behavior analysis. Adult flies were starved overnight (16 hours) and then fed a 5% sucrose solution containing DMSO or 60uM taxol for 5 hours prior to behavioral analysis.

### Electrophysiology

We adapted the methods of Gong et al (2013) to record NOMPC-mediated mechanically evoked responses from central neurons in the *Drosophila* ventral nerve cord (VNC). Adult physiology preparations were as previously described with some modification (Tuthill and Wilson, 2016). Flies were cold-anesthetized and fixed with their ventral side facing up to the underside of a custom-milled steel platform using UV-cured glue (KOA 300, KEMXERT). The ventral head and anterior thorax were partly inserted through a hole in the platform. The top side of the platform, and thus also the exposed parts of the head and thorax, were continually perfused with oxygenated saline. All six legs were glued to the holder with UV-cured glue. A small hole was manually dissected in the cuticle of the ventral thorax to expose the prothoracic neuromeres, and the perineural sheath was gently removed with fine forceps to expose neuronal cell bodies.

The preparation was perfused at ∼2-3 ml/min with saline (103 mM NaCl, 3 mM KCl, 5 mM TES, 8 mM trehalose, 10 mM glucose, 26 mM NaHCO_3_, 1 mM NaH_2_PO_4_, 1.5 mM CaCl_2_, and 4 mM MgCl_2_; pH 7.1, osmolality adjusted to 270-275 mOsm) bubbled with 95% O2/5% CO_2_. Recordings were performed at room temperature. Cell bodies were visualized using an infrared LED (Smartvision) and a 40× water-immersion objective on an upright compound microscope equipped with a fluorescence attachment (Sutter SOM). Whole-cell patch-clamp recordings were targeted to GFP-labeled cell bodies in the prothoracic region of the VNC. The internal patch pipette solution contained (in mM): 140 potassium aspartate, 10 HEPES, 1 EGTA, 4 MgATP, 0.5 Na_3_GTP, 1 KCl, and 13 biocytin hydrazide (pH 7.2, osmolarity adjusted to ∼265 mOsm). We distinguished motor neurons from other glutamatergic neurons by the characteristic and reliable positions of their cell bodies (Baek and Mann, 2009; Brierley et al., 2012), as well as their intrinsic properties (input resistance, resting membrane potential, and spike waveform).

All recordings were made in current-clamp mode using an Axopatch 700A amplifier. Data were low-pass filtered at 5 kHz before they were digitized at 10 kHz by a 16 bit A/D converter (National Instruments, USB-6343), and acquired in Matlab. Stable recordings were typically maintained for ∼1 hour. A small hyperpolarizing current (approximately -5 to -10 pA) was injected to compensate for the depolarizing seal conductance (Gouwens and Wilson, 2009). Analysis of electrophysiology data was performed with custom scripts written in MATLAB and Python.

Motor neurons were mechanically stimulated with a closed-loop piezoelectric actuator (Physik Instrumente P-841.60, 90 μm travel range, with E-509.S1 sensor/piezo servo-control module). Mechanical stimuli were generated in Matlab and sent to the amplifier at 5 kHz using an analog output DAQ (National Instruments 9263). Mechanical stimuli were generated in Matlab and sent to the amplifier at 5 kHz using an analog output DAQ (National Instruments 9263). The stimulating pipette was positioned next to the ipsilateral VNC neuromere under visual control. A 12 μm square wave was used to indent the neuropil. Because the motor neuron cell bodies are segregated from the VNC neuropil, the mechanical stimuli had no visible effect on the cell body and patch pipette. Stimulation of the contralateral neuromere failed to evoke a response (data not shown).

## Statistical Analysis

Datasets were tested for normality using Shapiro-Wilks goodness of fit tests. Details on statistical tests are provided in figure legends.

## Acknowledgements

This work was supported by a grant from the National Institutes of Health (NINDS R01 NS076614), a UW Research Innovation award, and startup funds from UW (J.Z.P.); support from Mark Peifer and a UNC Biology/iBGS Pilot Award (S.L.R); and by grants from the Human Frontiers in Science Program (RGY0090/2014) and National Institutes of Health (NINDS R01NS089787) (X.Y.). Fly Stocks obtained from the Bloomington *Drosophila* Stock Center (NIH P40OD018537) and antibodies from the Developmental Studies Hybridoma bank, created by the NICHD of the NIH and maintained at The University of Iowa, were used in this study. We thank Joshua Vaughan for assistance with Expansion microscopy, Mark Peifer and Peter Soba for critical reading of this manuscript.

## Figure Legends

**Figure S1.**
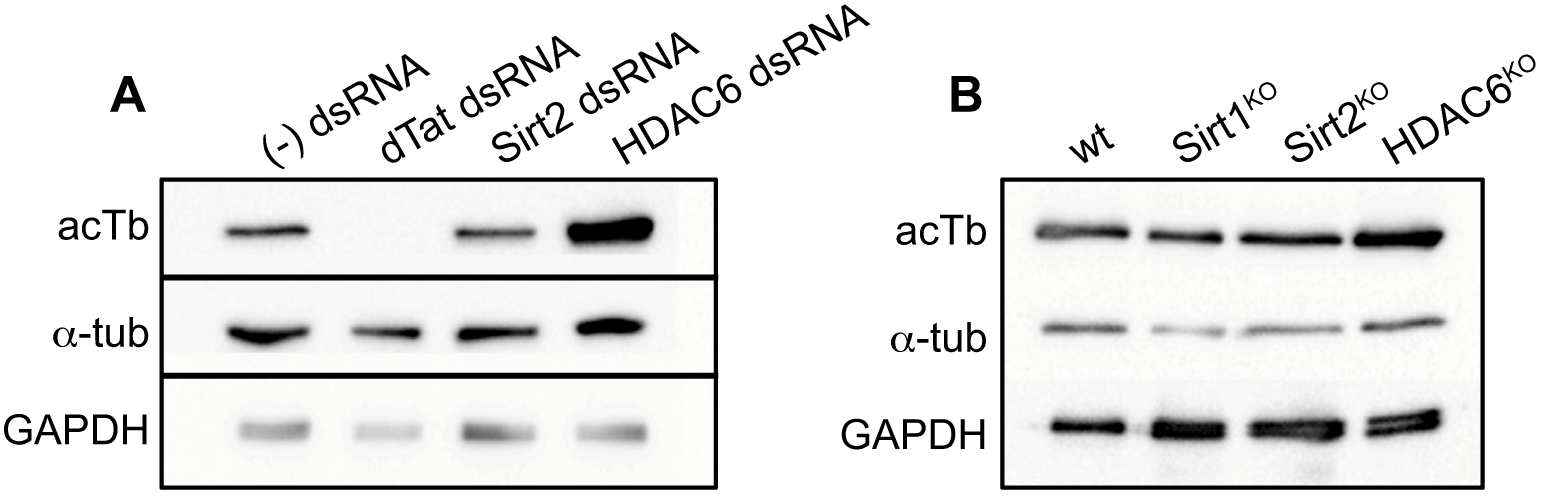
Related to Figure 1. (A) HDAC6 is the major α-tubulin deacetylase in *Drosophila.* (A) S2 cells were treated with dsRNA targeting dTat, Sirt2, HDAC6, or a negative control and the lysates were immunoblotted for acTb, total α-tubulin, and GAPDH as a loading control. (B) Lysates were prepared from 3rd instar larvae from wild type, *Sir2^KO^, Sirt2^KO^,* or *HDAC6^KO^* mutant fly lines and immunoblotted for acTb, total α-tubulin, and GAPDH as a loading control.

**Figure S2.**
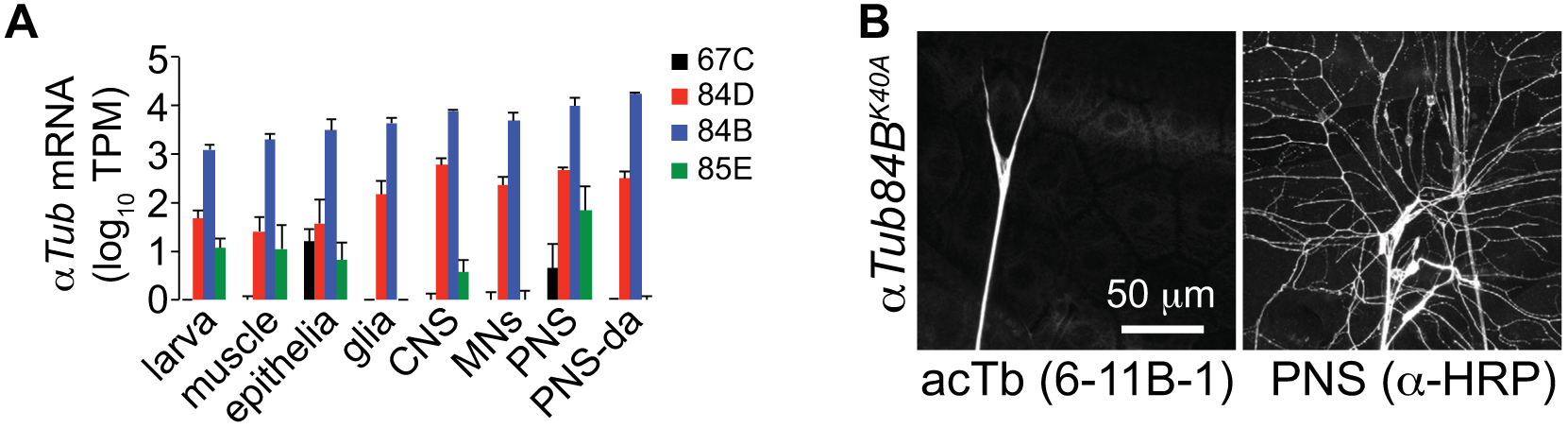
Related to Figure 3. αTub K40 is required for tubulin acetylation. (A) Larval mRNA expression of the *Drosophila α-tubulin* isotypes in the indicated cell types; *αTub84B* accounts for >90% of *α-tubulin* mRNA in the PNS. Reads were aligned to the *D. melanogaster* genome, FlyBase genome release 6.04 using TopHat2 version 2.0.14 and Bowtie2 version 2.2.3. (B) acTb immunoreactivity is absent in *αTub84B^K40A^* mutant third instar larva, demonstrating that the *αTub84B* isotype accounts for the majority of acTb in larvae.

**Figure S3.**
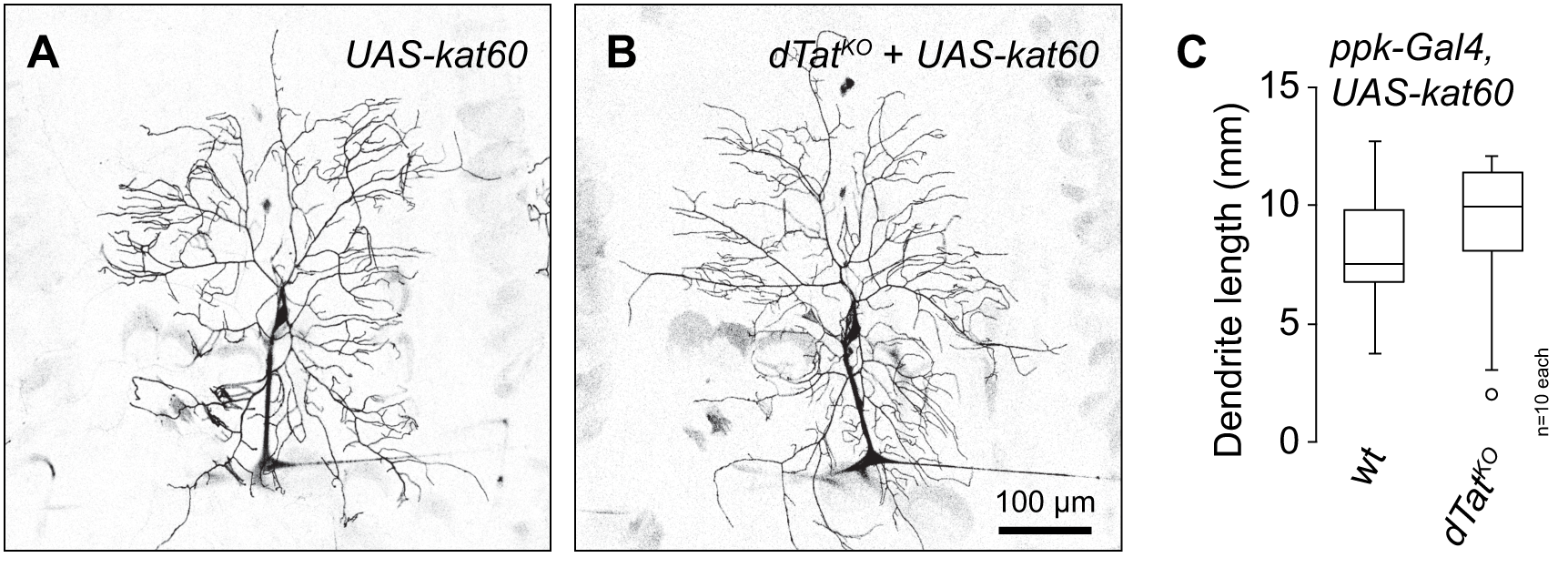
Related to Figure 4. *dTat* has no effect on *katanin*-induced remodeling of c4da dendrite arbors. Representative images of control larvae (A) or *dTat^KO^* mutant larvae (B) overexpressing *katanin (UAS-kat60)* in c4da neurons under the control of *ppk-Gal4.* Dendrites were labeled with the c4da-specific marker *ppk-CD4-tdTomato* visualized via live confocal imaging. (C) Quantification of total dendrite length for the indicated genotypes. Boxes mark 1st and 3rd quartiles, bands mark medians, whiskers mark 1.5 x IQR, and outliers are shown as points. Welch’s test of unequal variance revealed no significant difference was detected between the two groups.

**Figure S4.**
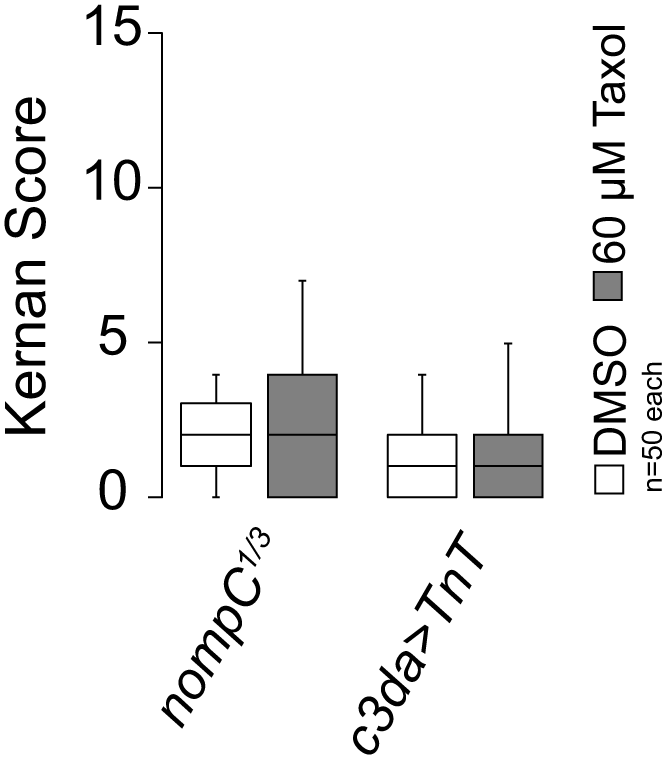
Related to Figure 7. Taxol enhancement of gentle touch responses depends on activity of NOMPC and c3da neurons. Boxplots depict behavioral responses of third instar larvae of the indicated gentoypes to gentle touch stimulus at 96 h after egg laying (AEL). Mutation of *nompC* or blocking synaptic transmission by expressing Tetanus toxin in c3da neurons *(c3da>TnT)* rendered taxol-fed larvae insensitive to gentle touch.

**Table S1.** Related to Figure S5. Expression levels of mechanosensory channels in different cell types.

|  |  |  |  |
| --- | --- | --- | --- |
|  | PNS | da neurons | motoneurons |
| *CG46121* | 56.91 | 7.10 | 0 |
| *iav* | 3.26 | 0 | 4.59 |
| *nan* | 0 | 0.02 | 0.19 |
| *nompC* | 87.84 | 22.92 | 0 |
| *pain* | 6.23 | 4.08 | 0.51 |
| *Piezo* | 16.93 | 100.48 | 4.00 |
| *ppk* | 272.00 | 398.80 | 0 |
| *ppk26* | 628.29 | 650.44 | 0 |
| *rpk* | 0 | 0 | 0 |
| *trp* | 0 | 0 | 0 |
| *Trpgamma* | 0 | 0 | 0.02 |
| *trpl* | 20.83 | 0.03 | 0 |
| *wtrw* | 0.25 | 0 | 0 |

Mean mRNA expression levels (tpm, transcripts per million) are shown for mechanosensory channel genes in each of the indicated cell types. N = 4 independent samples for PNS and da neurons; 7 independent samples for motoneurons.

**Table S2.**
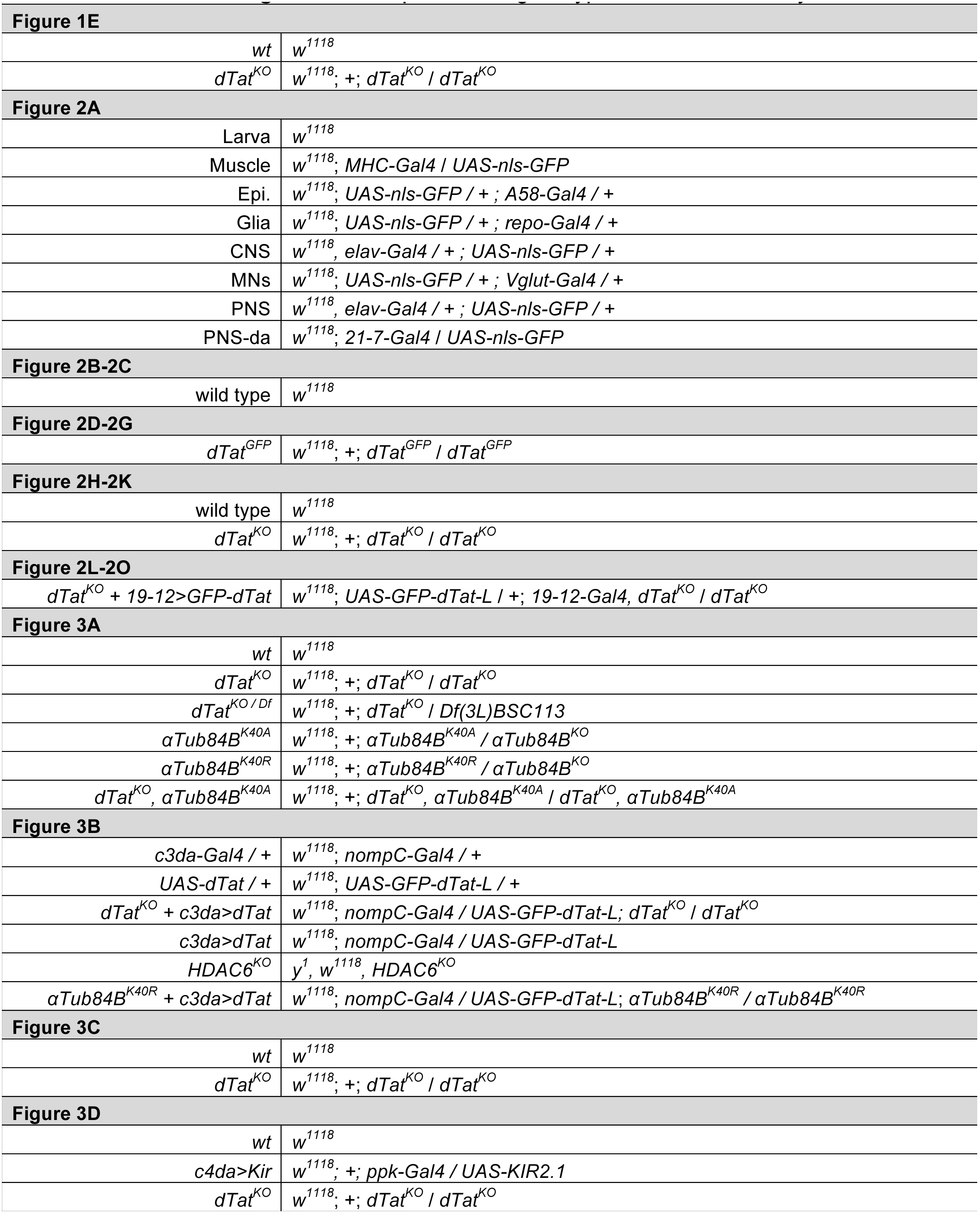
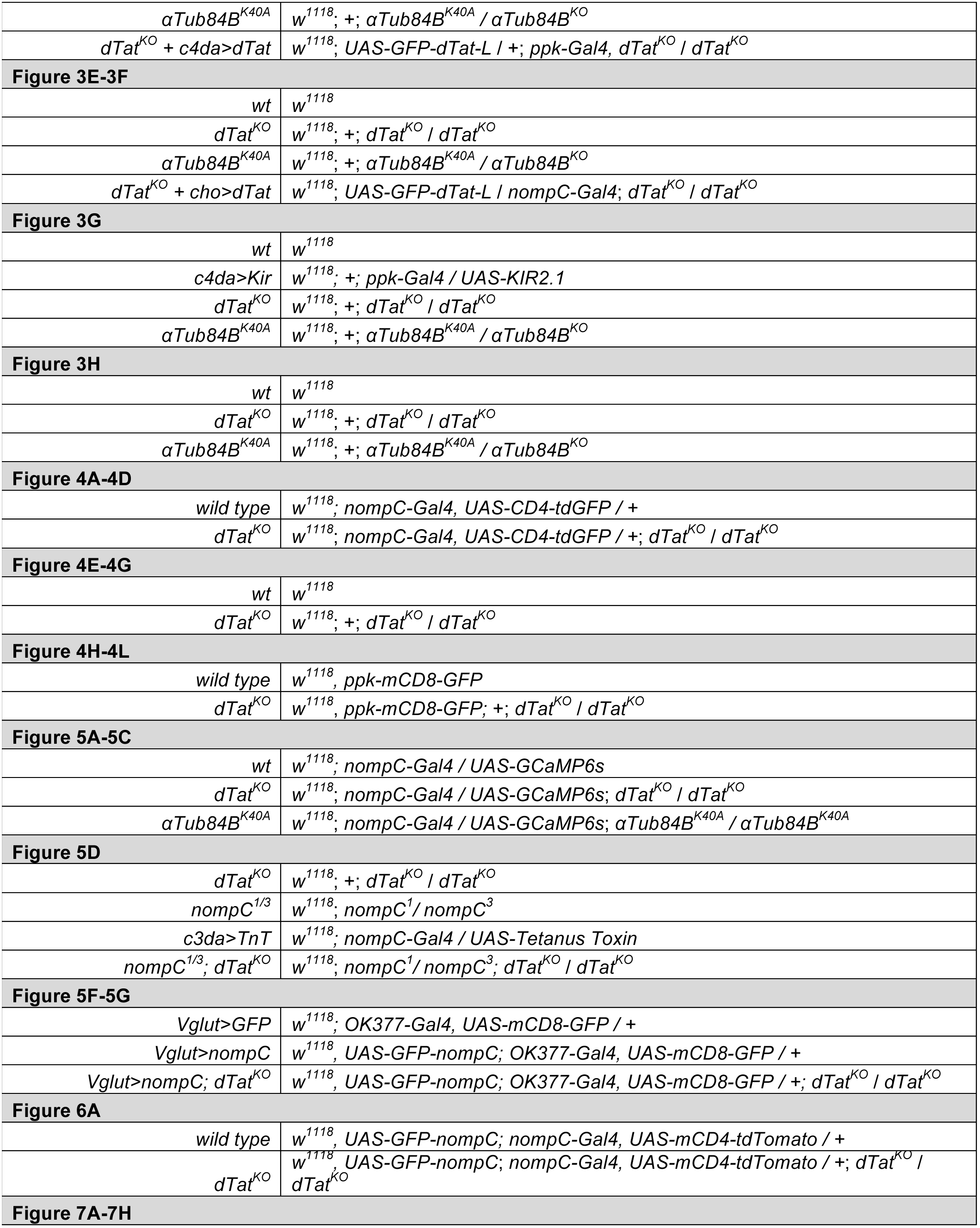
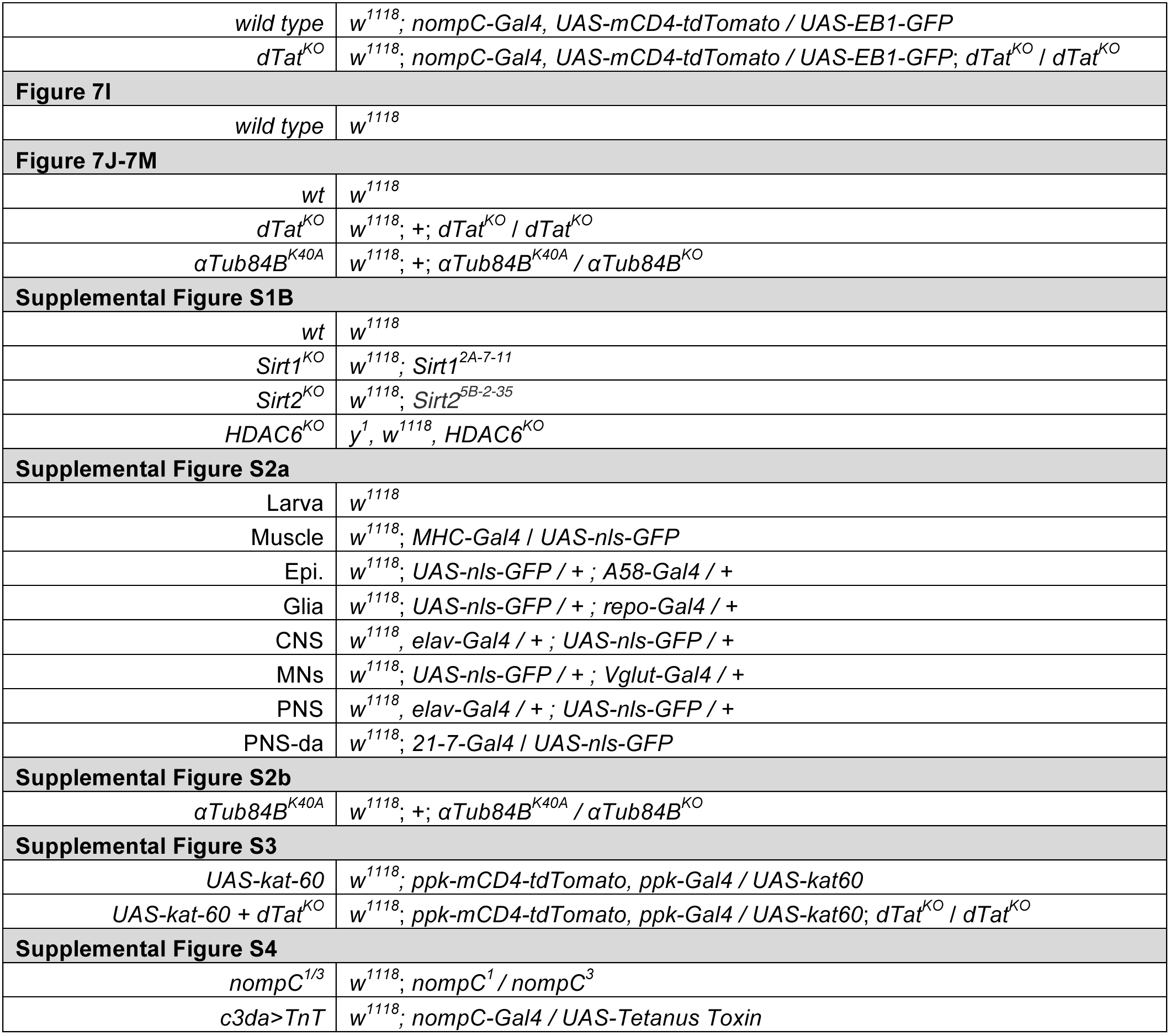
Related to Figures 1-7. Experimental genotypes used in this study.

**Table S3.**
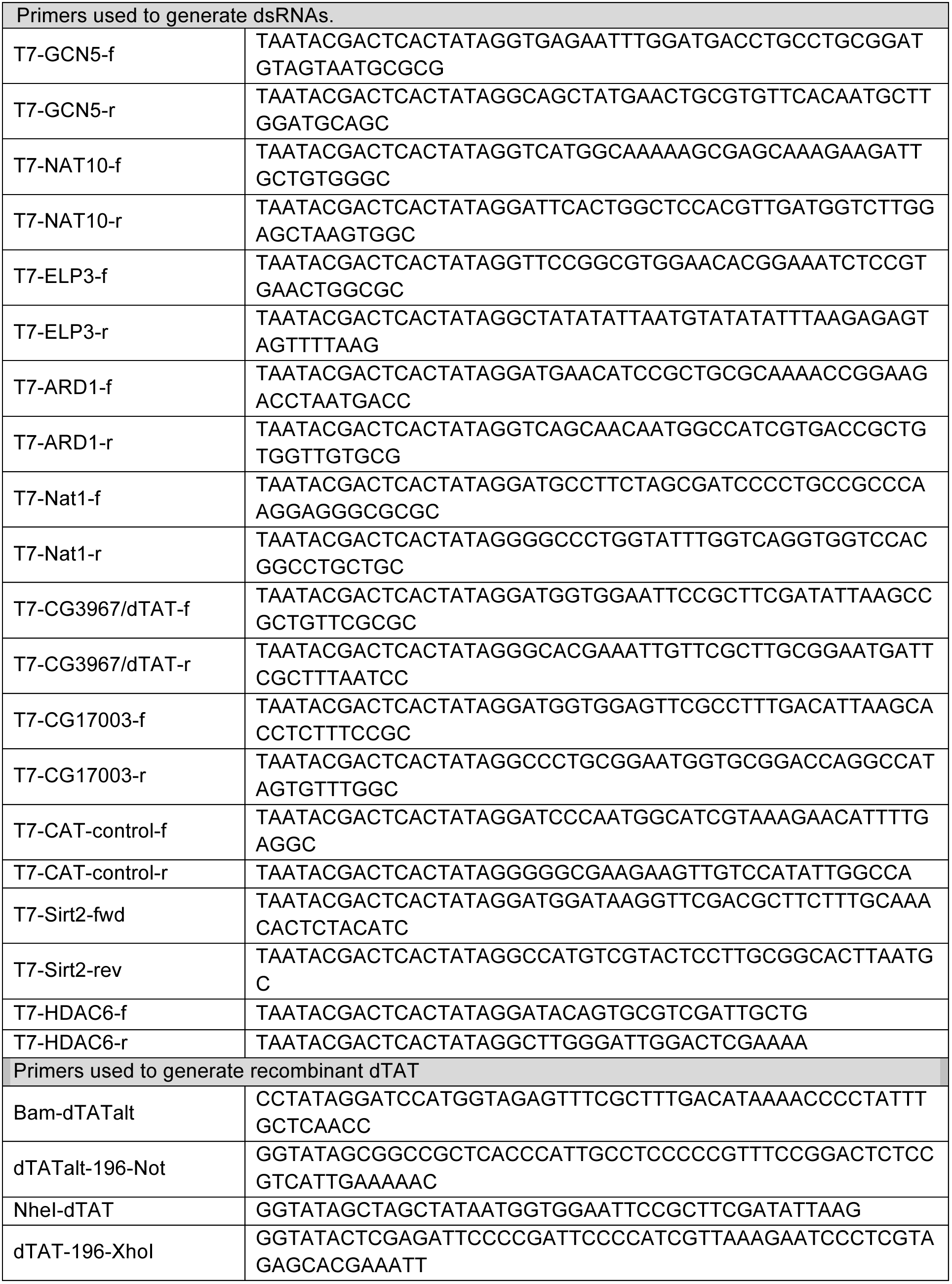
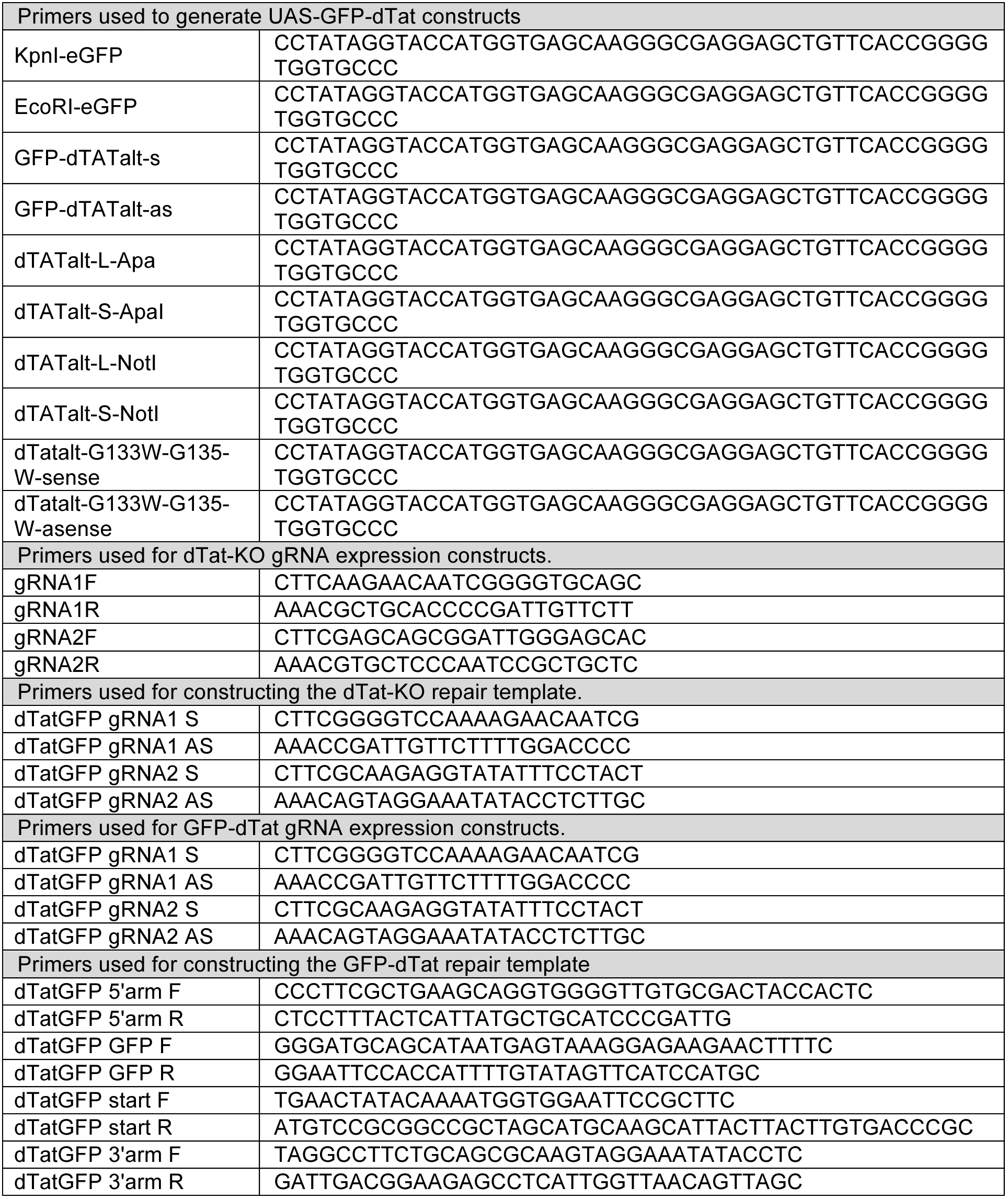
Oligonucleotide primers used in this study.

## Abbreviations

acTb: acetylated tubulin
AR: ankyrin repeats
K40: lysine 40 of α-tubulin
PNS: peripheral nervous system
PTM: post-translational modification
TRP: transient receptor potential channel

